# Serpin-Driven Green Camouflage and NIR Fluorescence in Frogs

**DOI:** 10.64898/2026.02.11.704363

**Authors:** Ravichandran Vignesh, Tri Vu, Grace Harvey, Luca Menozzi, Jesse Delia, William White, Pohan Lin, Erini Galatis, Sönke Johnsen, Junjie Yao, Carlos Taboada

## Abstract

Animals have evolved multiple strategies to generate optical traits and coloration. While most amphibians rely on a three-dimensional arrangement of chromatophores in the skin, hundreds of arboreal frog species achieve leaf-like green color through a different mechanism involving reduced pigmentation, subcutaneous biological mirrors, and high concentrations of the blood-derived pigment biliverdin. Although biliverdin is rapidly excreted in most vertebrates, hylid and centrolenid frogs can retain it through the binding to a serpin-family protein (BBS). Here we show that BBSs bind biliverdin with high affinity (K_d_ < 10 nM), comparable to hormone–receptor interactions. This interaction alters biliverdin’s spectral signature in ways that resemble those of green-leaf pigments. BBSs from different species exhibit distinct biophysical properties, accounting for interspecific differences in color saturation and hue. Unlike most serpins *in vivo*, BBS of the glassfrog *Teratohyla pulverata* is naturally cleaved, yielding a highly thermostable, thermodynamically favored protein, without loss of affinity. Using custom-designed hyperspectral photoacoustic tomography (PAT), we demonstrate that BBS is widely distributed throughout the body, contributing to whole-body color and camouflage. Furthermore, we show that BBSs emit near-infrared (NIR) fluorescence (>700 nm) rendering these frogs fluorescent in a spectral region where biological tissues are largely transparent. Together, BBSs shed light on serpin evolution, protein thermostability, amphibian color diversity, and the development of NIR molecular probes.

## Introduction

In most ectothermic vertebrates, coloration is primarily produced by skin chromatophores^1–3^. However, multiple arboreal frog lineages have evolved an alternative mechanism characterized by reduced chromatophores and the high accumulation of the bile pigment biliverdin throughout their tissues –including blood, lymph, muscle, and bones^4–6^. This accumulation of biliverdin, a condition known as chlorosis, modulates light absorption and transmission resulting in leaf-like green camouflage^5^.

Biliverdin is the first breakdown product of heme from senescent red blood cells and exhibits green coloration due to absorption in the violet (Soret band) and red (Q band) regions of the spectrum^7^. Although intrinsically green, biliverdin’s absorption spectrum can be strongly influenced by its conformation, protonation state, and interactions with its molecular environment^7,8^. Leaf-dwelling frogs in the families Hylidae and Centrolenidae have independently co-opted a liver-derived serpin, termed the biliverdin-binding serpin (BBS), which binds biliverdin and exploits its spectral tunability^5,9^. By altering biliverdin’s conformation and electronic structure, BBSs shift both the position and magnitude of its absorption peaks toward that approximate those of plant chlorophyll^5,9^. These spectral changes include a red shift of the Soret band and a more structured, blue-shifted Q band that exhibits a threefold increase in absorption^5^. Across frog lineages, amino acid substitutions in BBSs produced distinct absorption profiles that shaped frog coloration, resulting in optical phenotypes ranging from bluish to light green^5,9^.

Biliverdin-binding in proteins is typically associated with a rigidification of biliverdin, which results in fluorescence emission, primarily in the red (630–700 nm) and near-infrared (NIR, >700 nm) regions of the spectrum^10–15^. This photochemical behavior has important biological and practical implications. The NIR region corresponds to the optical transparency window, where light absorption and scattering by animal tissues are minimal^16–18^. This suggests that frogs expressing BBSs could emit NIR fluorescence, even from deep tissues *in vivo*—a characteristic shared with chlorophyll-bearing vegetation^19–21^ that may expand the repertoire of optical traits in amphibians and enhance their leaf-like camouflage. Whether BBSs indeed produce detectable fluorescence *in vivo* remains unknown, as quenchers present in frog tissues under normal physiological conditions could attenuate BBS fluorescence.

The use of biliverdin to create colors in vertebrates presents physiological challenges due to its rapid excretion or enzymatic reduction to bilirubin^1,22–26^, even under extreme hemolysis or following direct biliverdin administration^6,23,27–29^. This physiological constraint has biological consequences, as biliverdin is one of the few green pigments in animals, yet its use to modulate colors in nature is rare^30,31^. Furthermore, for practical applications, the use of biliverdin-binding proteins that rely on endogenous biliverdin is often impaired by the short-lived nature of biliverdin in tissues^13,32,33^. Preserving high concentrations of biliverdin *in vivo* thus requires mechanisms to minimize its chemical degradation and to slow its natural clearance^32,34^. The widespread use of BBSs among leaf-dwelling frogs of different lineages suggests that BBSs have convergently evolved mechanisms to bind biliverdin with high affinity.

Understanding the biophysical properties that enable biliverdin binding and spectral tuning in BBSs will shed light on the evolution of protein-based coloration and provide a framework for linking the molecular evolution of a single protein to a leaf-like optical phenotype. Additionally, their fluorescence in the NIR positions BBSs as a promising template in developing bioinspired optical probes that permit deep tissue penetration. Finally, identifying BBS variants that have been evolutionarily tuned to enhance biliverdin binding may reveal molecular strategies for overcoming the rapid turnover of biliverdin *in vivo*—a key limitation of NIR fluorescent probes that rely on endogenous biliverdin^13,32,35–37^.

Here, we describe a new biliverdin-binding serpin (TpBBS) from the glassfrog *Teratohyla pulverata* (Centrolenidae) and characterize its biophysical and optical properties. We show that TpBBS is a hyperstable protein that binds biliverdin with high affinity, approaching that of hormone–carrier and antibody–epitope interactions^38–46^. Extending our analysis to two additional independent origins of BBSs in treefrogs (*Boana punctata* and *Sphaenorhynchus lacteus*), we demonstrate that distinct BBSs adopt different conformational states while retaining strong biliverdin binding and producing species-specific green coloration. Using spectroscopy, NIR fluorescence imaging, and hyperspectral photoacoustic tomography, we further show that BBSs preserve their optical properties *in vivo*, substantially extending the known spectral range of amphibian fluorescence.

## Results

### BBSs and green leaf-color camouflage

The arboreal frogs *Teratohyla pulverata*, *Boana punctata*, and *Sphaenorhynchus lacteus* examined in this study (**Fig. 1**) represent three independent evolutionary origins of BBSs^5^. Their translucent skin reveals large accumulations of subcutaneous blue lymph. To investigate the role of BBSs in producing plant-like green coloration, we purified BBSs from each species—TpBBS, BpBBS, and SpBBS (**Fig. 2A**)—and characterized their biophysical and optical properties.

**Figure 1.**
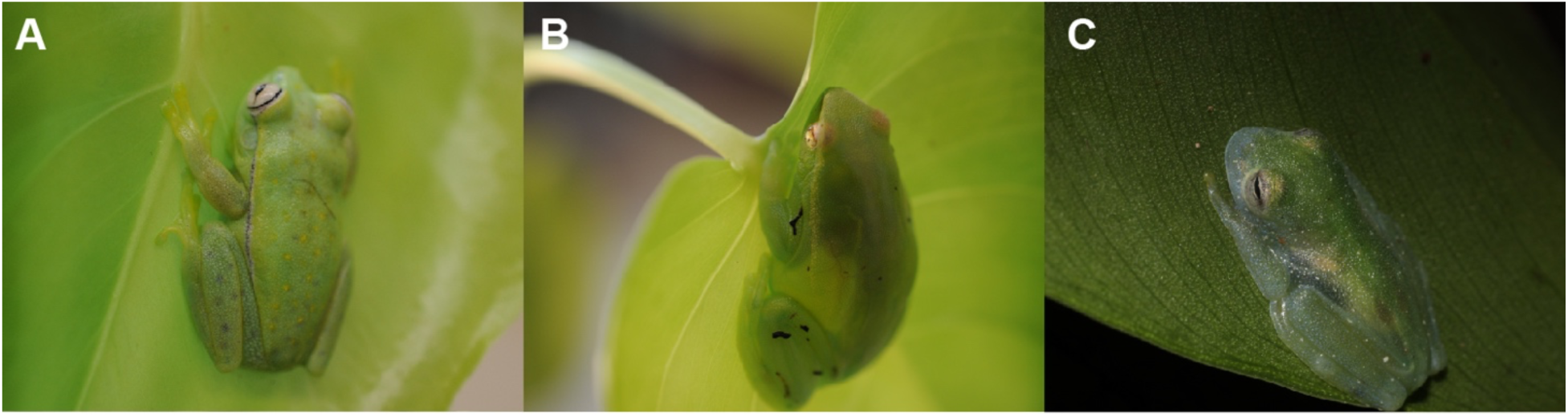
Chlorotic arboreal frogs of the Hylidae and Centrolenidae family. **(A)** The polka-dot treefrog *Boana punctata* from Suriname, **(B)** the hatchet-faced treefrog *Sphaenorhynchus lacteus* from Suriname, and **(C)** the powdered glassfrog *Teratohyla pulverata* from Nicaragua. These three species represent three independent evolutionary origins of high BBSs expression, tissue transparency, and biological mirrors coating one or more organs or parietal peritonea.

**Figure 2.**
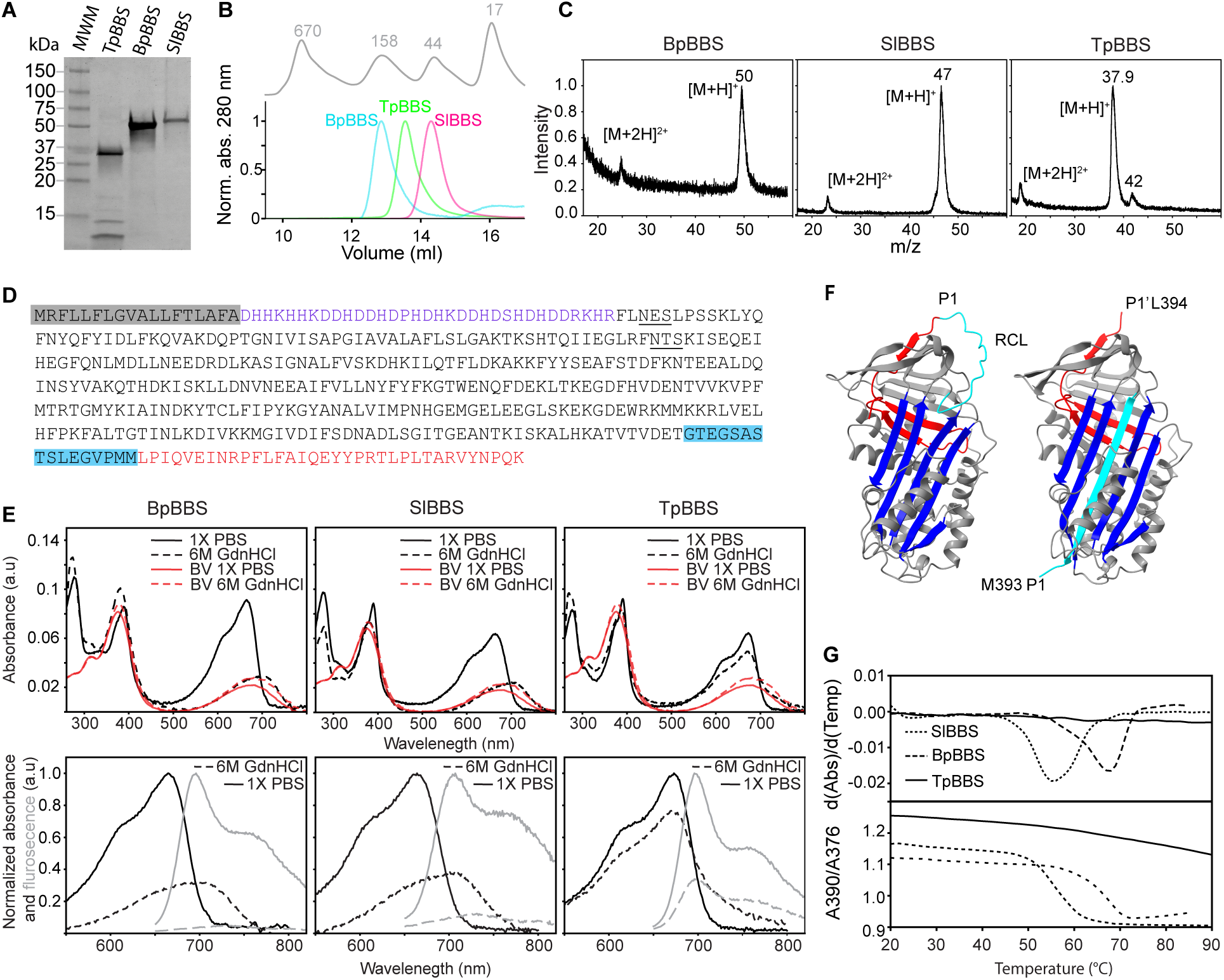
Purification and characterization of BBSs from chlorotic frogs. **(A)** Coomassie Brilliant Blue-stained SDS-PAGE of purified BBS from the tissues of *B. punctata*, *S. lacteus*, and *T. pulverata.* **(B)** Size Exclusion Chromatogram of purified endogenous BpBBS (cyan), TpBBS (green), SlBBS (pink), and standards (gray). **(C)** MALDI-TOF of the different BBSs. **(D)** Amino acid sequence of TpBBS identified from *de novo* transcriptomics and proteomics; signal peptide (gray), N-terminus (purple), RCL (cyan), C-terminus (red), with potential glycosylation sites underlined. **(E)** *Top:* Absorbance spectra for Bp-, Sl-, and TpBBSs in 1X PBS and denaturing conditions (6M GdnHCl) compared to free biliverdin (BV). *Bottom*: Normalized Q band absorbance (black) and fluorescence (gray) of BBSs in 1X PBS (solid lines) and in 6 M GdnHCl (dotted line) respectively. **(F)** AlphaFold predicted structure of TpBBS, uncleaved stressed confirmation (*left*) and hyperstable relaxed conformation, with the RCL inserted into β-sheet A (cyan) of the serpin fold (*right*). **(G)** Melting curves of different BBS (Absorbance vs T) representing the loss of structural fold and release of biliverdin (*bottom*) and the first derivative (dA/dT vs T) (*top*).

Size exclusion analysis of the purified endogenous proteins revealed a symmetric peak for all the BBSs, with the predicted hydrodynamic radii ordered as BpBBS>TpBBS>SlBBS (**Fig. 2B**). The molecular weights of the purified BBSs were estimated by MALDI-TOF to be 37.9 kDa for TpBBS, 47 kDa for SlBBS, and 50 kDa for BpBBS (**Fig. 2C**).

Although the sequences of BpBBS and SlBBS have been previously reported^5^, and we determined the full sequence of TpBBS using tandem mass spectrometry, searching the resulting peptides against the *de novo* liver transcriptome of *T. pulverata* (**Fig. 2D**). The predicted molecular weight of TpBBS is 46.5 kDa, and the sequence includes two consensus N-glycosylation motifs (**Fig. 2D**). Multiple sequence alignment of TpBBS, SlBBS, BpBBS (**Supp.Fig.1A**) and other serpins shows that TpBBS shares 76–83% sequence identity with other glassfrogs (Centrolenidae), but only 46–52% identity with treefrogs of the Hylidae family and serpins (**Supp. Fig. 1B**).

Absorbance and fluorescence spectra for all BBSs are shown in **Fig. 2E**. The Soret band exhibited maximal absorbance at 389–392 nm across all BBSs, red-shifted relative to free biliverdin in 1X PBS (**Fig. 2E**). The Q band, in contrast, varied in both spectral shape and peak position; TpBBS showed a maximum at 673–675 nm, whereas BpBBS and SlBBS peaked at 666 nm (**Fig. 2E**). Upon excitation at either 390 or 630 nm, all BBSs showed a broad fluorescence emission with a peak at 693–707 nm, extending into the near-infrared region of the spectrum (**Fig. 2E**).

BpBBS is known to contain two N-glycosylation sites^5^, whereas SlBBS possesses a single glycosylation consensus sequence. To assess glycosylation as a post-translational modification in the different BBSs, we treated the samples with PNGase F. BpBBS and SlBBS were deglycosylated upon PNGase F treatment (**Supp. Fig. 2A**). In contrast, the molecular weight of TpBBS remained unchanged after PNGase F treatment (**Supp. Fig. 2A**), suggesting that it is either not glycosylated, carries glycans of low molecular weight, or contains glycans resistant to PNGase F cleavage.

### TpBBS is a thermally stable biliverdin-binding serpin in the relaxed conformation

Most serpins are serine protease inhibitors that regulate protease activity in diverse physiological processes employing a unique mechanism centered on a structural feature called the reactive center loop (RCL)^47^. Inhibition occurs when a protease cleaves the RCL, which drives the conformational change causing the insertion of RCL into the central β-sheet A of the serpin fold^48^ (**Fig 2F**). This dramatic structural transition is possible because the native fold is metastable, or “*stressed*”, with the exposed RCL stable under physiological conditions, yet energetically far from the protein’s optimal state^49^. After RCL is cleaved and inserted into the serpin fold, it adopts a “*relaxed*” conformation that enhances serpin stability, reflected in an increased melting temperature (T_m_)^50^. While the empirical molecular weights of BpBBS and SlBBS matched their theoretical predictions, TpBBS appeared smaller in both SDS PAGE and MALDI-TOF, suggesting that in *T. pulverata*, TpBBS might be naturally cleaved. Bottom-up mass spectrometry of the purified TpBBS gel band (**Fig. 2A, C**), revealed ∼80% sequence coverage and did not include any peptides from either the N- or C-terminal regions (**Fig. 2D**), suggesting that TpBBS is truncated at both ends. To determine whether the cleaved fragment is associated with the protein in the native state, we crosslinked TpBBS (**Supp Fig. 2B**) and analyzed the sample with MS/MS after trypsinization. The crosslinked sample contained the C-terminal peptide (**Supp. Fig. 2C**), bringing the coverage to 91% (**Supp. Fig. 2D**), but showed no evidence of the missing N-terminal region, indicating that this portion is either absent or extensively modified. The cleavage in TpBBS was further confirmed by mass spectrometry analysis of the native TpBBS, which revealed an additional peak at 4,088 m/z (**Supp. Fig. 2E**). Analysis of the fragmentation patterns of this peptide indicated that TpBBS is naturally cleaved at M393 (**Supp Fig. 2C-E**) in its native state, within the RCL (**Supp. Fig. 1A**), which suggests that the protein might adopt a hyperstable relaxed conformation, with the RCL inserted into β-sheet A of the serpin fold (**Fig. 2F**). To test this hypothesis—and to compare it with BpBBS and SlBBS, which are presumably in the native stressed conformation—we incubated all three proteins in 6 M guanidine hydrochloride (GdnHCl) for one hour. In BpBBS and SlBBS, biliverdin rapidly dissociated from the apoprotein upon denaturation, as evidenced by a 2–3-fold decrease in Q band absorbance (**Fig. 2E**), a hypsochromic shift of the Soret band from 390 to 376 nm, and a pronounced reduction in near-infrared fluorescence emission (**Fig. 2E**). In contrast, TpBBS exhibited only minor changes in absorbance and NIR fluorescence upon GdnHCl treatment, indicating a highly stable fold typical of serpins in the relaxed conformation (**Fig. 2F**)^48,49^.

To further assess whether the endogenous TpBBS is in relaxed conformation, we performed a thermal unfolding of all the BBSs and calculated their melting temperature (T_m_). The melting curves of BBS (A390/A376 *vs* T) and the first derivative (dA/dT *vs* T) are shown in **Fig. 2G**. T_m_ values for BpBBS and SlBBS were calculated to be 67 °C and 56 °C, respectively, whereas TpBBS showed no temperature-dependent change in the ratio of absorbance. These results indicate that TpBBS is highly stable, consistent with the behavior observed under GdnHCl denaturation. This exceptional stability supports the notion that TpBBS exists in the thermodynamically more stable “relaxed” conformation, with the cleaved RCL inserted into β-sheet A (**Fig. 2F**).

### BBSs bind biliverdin with high affinity

Chlorotic frogs can accumulate high concentrations of biliverdin, which most organisms would rapidly excrete or enzymatically reduce to bilirubin^25,28,33^. This suggests their BBSs possess high affinity for biliverdin, thereby extending its lifetime *in vivo* by sequestration and preventing excretion^4,5^. To evaluate the affinity of BBSs for biliverdin, we expressed recombinant BBSs in *E. coli* and performed fluorescence titration experiments in the near-infrared (NIR) range. To verify whether the recombinant proteins retained biliverdin binding properties comparable to their endogenous counterparts, we recorded absorbance and fluorescence spectra for both sets of proteins (**Fig. 3A-C**). The spectra were indistinguishable in both the Soret and Q bands, in terms of amplitude and peak position, indicating identical optical properties. The extinction coefficients of the Soret and Q band were determined for all three biliverdin-bound BBSs (**Table 1**) and revealed a 2.5–3.5 times increase in absorption in the Q band relative to free biliverdin (**Supp. Fig. 3**). The change in optical properties induced by the binding of biliverdin to recombinant BBSs is observed as an increase in the color saturation of the solution and the emergence of fluorescence emission upon excitation in the far-red region (**Fig. 3D–F**). The quantum yield of the proteins upon excitation at 650 nm was 1.39% for BpBBS, 0.58% for SlBBS, and 1.24% for TpBBS.

**Fig. 3.**
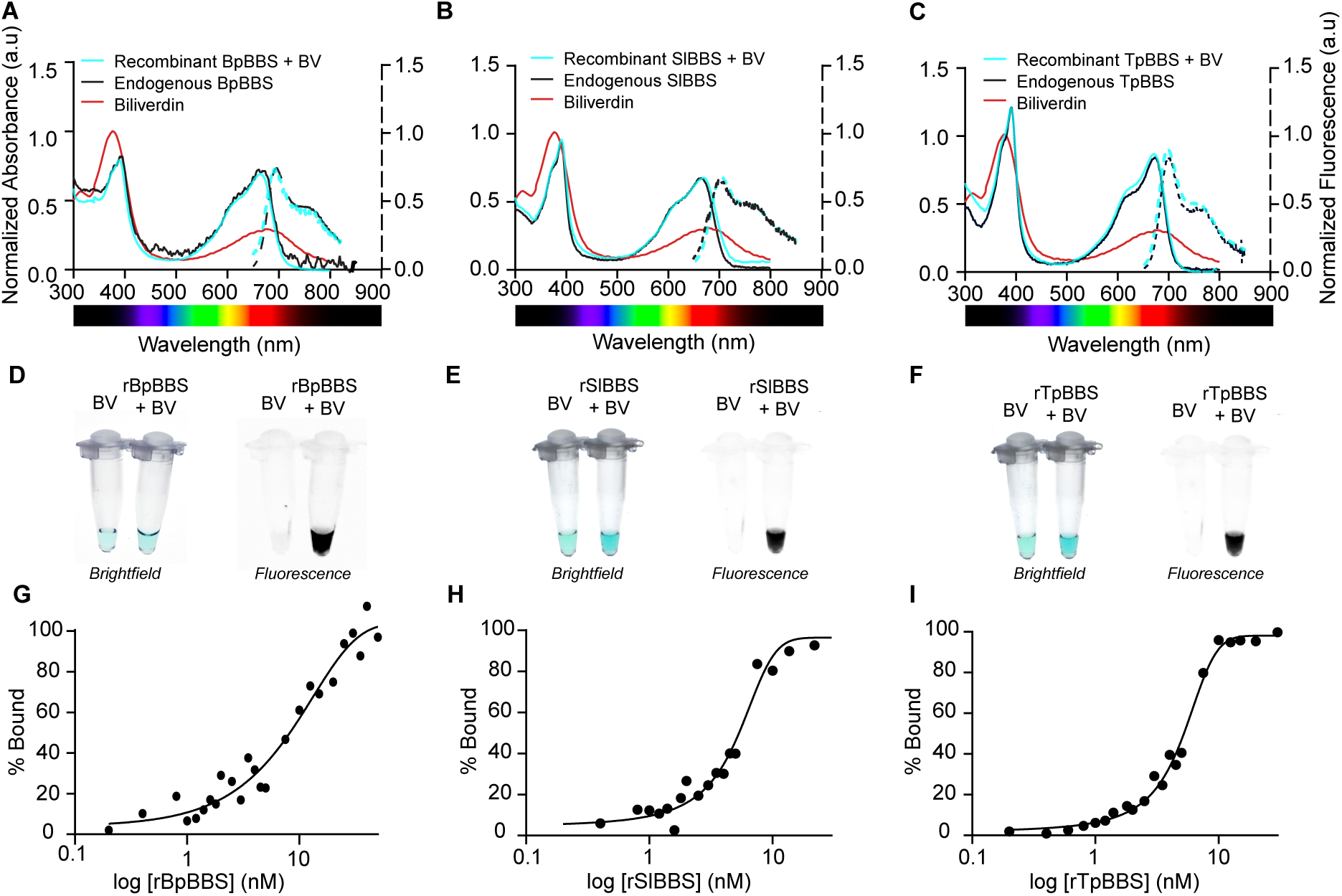
Biliverdin binding in the recombinant BBS. (A–C) Normalized absorbance (normalized to the Soret band of free biliverdin (BV) at the same concentration) and fluorescence (dotted lines, excitation 630 nm for BpBBS, and 390 nm for SlBBS and TpBBS, emission 650 nm-850 nm) of biliverdin-bound recombinant BBSs (cyan) compared to endogenous BBSs (black) and free biliverdin (red). **(D–F)** Brightfield and fluorescence photos of free biliverdin and recombinant biliverdin-bound BBSs. The color of biliverdin changes from green to blue/blueish-green, and samples become fluorescent. Fluorescence images were taken on the Bio-Rad ChemidocMP Imager under Cy5.5 filter. **(G–I)** Ligand binding curves (%bound *vs* concentration) of recombinant BBSs after 4 h of equilibration, obtained by titrating biliverdin with increasing concentrations of recombinant apoprotein (0.02 nM-30 nM) to biliverdin (15 nM). Apparent K_d_ values: 7.5 nM for BpBBS, 5 nM for SlBBS, and 5 nM for TpBBS.

**Table 1.** Optical and binding properties of recombinant BBSs.

| Sample | $\epsilon_{280}$<br>($M^{-1}cm^{-1}$) | $\epsilon_{Soret}$<br>($M^{-1}cm^{-1}$) | $\epsilon_{Q-band}$<br>($M^{-1}cm^{-1}$) | $K_d$<br>(nM) | Quantum<br>Yield |
| --- | --- | --- | --- | --- | --- |
| BpBBS | 28,880 | 32,072 | 28,034 | 7.5 | 1.39% |
| SIBBS | 35,870 | 38,393 | 26,229 | 5.0 | 0.58% |
| TpBBS | 33,350 | 52,809 | 36,449 | 5.0 | 1.24% |

To quantify the interaction between BBS and biliverdin, we measured binding by keeping biliverdin at a fixed concentration (limiting reagent) while titrating with increasing amounts of recombinant apoprotein. We repeated the experiments at different biliverdin concentrations and equilibration times (**Supp. Fig. 4**) to ensure measurements were conducted in the binding rather than titration regime^51^. At biliverdin concentration as low as 15 nM, the apparent K_d_ values were 5 nM for rTpBBS, 5 nM for rSlBBS, and 7.5 nM for rBpBBS—still within the titration regime (estimated K_d_ ≈ ½ [limiting reagent]) (**Fig. 3G–I**). Further testing at lower protein concentrations was limited by the sensitivity of the fluorescence assay.

**Fig. 4.**
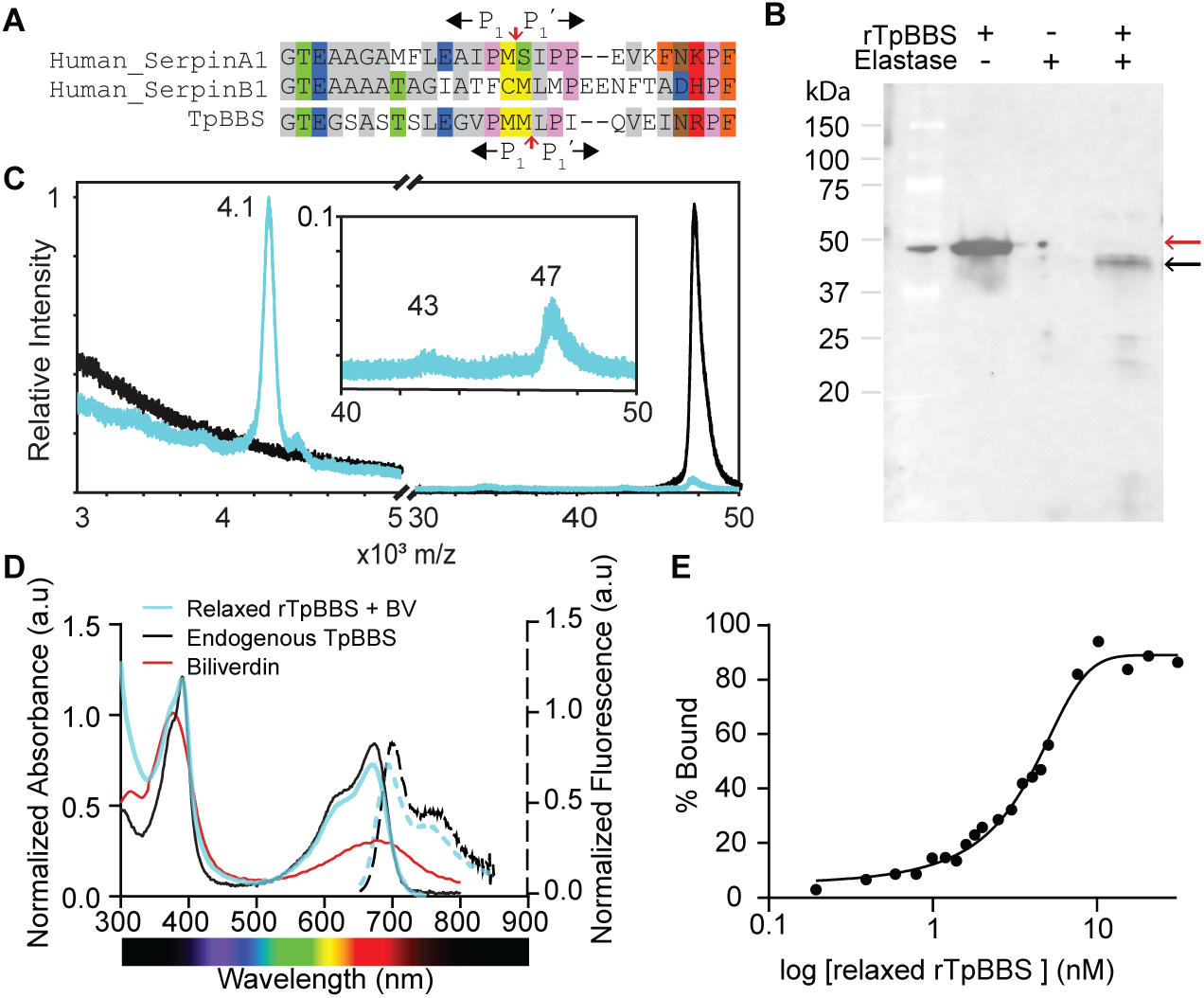
Relaxed rTpBBS binds biliverdin with high affinity. **(A)** Sequence alignment of RCL sequences of known neutrophil elastase inhibitors and TpBBS. The scissile bond of TpBBS is shifted one amino acid relative to the other canonical inhibitors (red arrow). **(B)** Cleavage of rTpBBS by (human) neutrophil elastase monitored with western blotting using anti-his antibody. The cleaved rTpBBS (black arrow) exhibited lower molecular weight than the uncleaved protein (red arrow). **(C)** Normalized MALDI-TOF of the relaxed rTpBBS (blue) compared to untreated rTpBBS (black). The peak at ∼4.1 kDa confirms the cleavage of rTpBBS at the scissile bond, in the same position as the endogenous TpBBS. *Inset,* shows the cleaved protein (43kDa) as analyzed by MALDI-TOF. **(D)** Absorbance spectra of relaxed rTpBBS bound to biliverdin (blue) normalized to the Soret band of free biliverdin at the same concentration, and fluorescence (dotted lines, excitation 390 nm, emission 650 nm-850 nm), endogenous TpBBS (black), and free biliverdin (red). **(E)** Ligand binding curves (%bound *vs* concentration) of relaxed rTpBBS after 4 h of equilibration. A fixed concentration of biliverdin (15 nM) was titrated with increasing concentrations of recombinant apoprotein (0.02 nM-30 nM). Apparent K_d_ of 3.5 nM for the relaxed rTpBBS.

### TpBBS has a high affinity for biliverdin in the stressed and relaxed state

While the protease recognition sites and inhibitory roles of specific serpins have been extensively characterized in humans and mice^48,49,52–58^, information on amphibian serpins remains limited^5,15,48^.

Given that endogenous TpBBS is cleaved in its relaxed serpin conformation—a structural state that, in other serpins such as the hormone carrier corticosteroid binding globulin (CBG) and others, has been associated with a drastic decrease in ligand affinity^59–63^—we decided to investigate whether the relaxation observed *in vivo* affects biliverdin binding.

To identify a protease capable of cleaving the TpBBS RCL for further biochemical tests, we aligned its sequence with those of human serpins (**Fig. 4A**). RCL of TpBBS shares the highest sequence identity with human SerpinA1 (α₁-antitrypsin), a well-characterized inhibitor of neutrophil elastase^47,52,64,65^. Hence, we predicted the presence of a human neutrophil elastase cleavage site in these BBSs (**Fig. 4A**). We, therefore, incubated the recombinant TpBBS (rTpBBS) with human neutrophile elastase and observed the proteolytic digestion, resulting in a reduction in apparent molecular weight, consistent with the loss of the C-terminal portion following cleavage at the scissile bond (**Fig. 4B**). Using MALDI-TOF, we confirmed that the released peptide is ∼4,08 Da (**Fig. 4C, Supp. Fig. 5**), consistent with the C-terminal fragment observed in the endogenous protein (**Supp. Fig. 2E**). These results indicate that the scissile bond in TpBBS is shifted by one position toward the C-terminus relative to other elastase inhibitors, such as SerpinA1 and Serpin B1 (**Fig. 4A**). Titration of the relaxed rTpBBS with biliverdin resulted in spectral features (absorbance and fluorescence) matching the biliverdin-bound endogenous TpBBS (**Fig. 4D**). The determined apparent K_d_ was 3.5 nM (**Fig. 4E**), which suggests that the high affinity for biliverdin is not impacted by the proteolytic cleavage of rTpBBS.

**Fig. 5.**
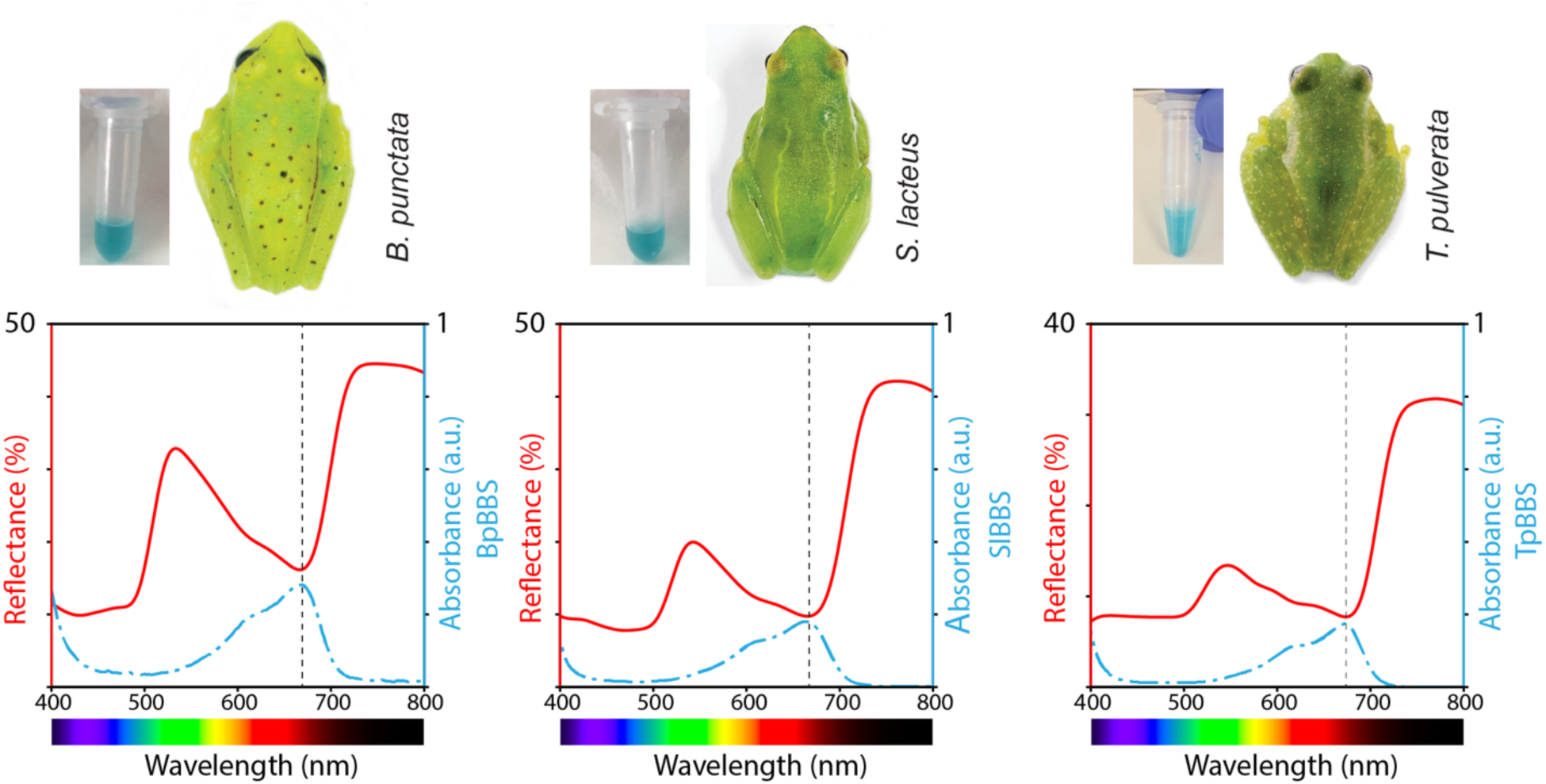
Whole-body reflectance and BBS absorbance in two treefrogs and a glassfrog. Reflectance spectra of chlorotic frogs (from left to right: *B. punctata*, *S. lacteus*, *T. pulverata*) show a peak in the green portion of the spectrum, a local minimum in the far-red, and a rapid increase of reflectance in the red-NIR boundary, known as red-edge that characterizes living, photosynthesizing vegetation^66^. The BBS used for absorbance measurements, extracted from each species, is shown next to the frogs. The peak and shape of BBSs’ absorbance perfectly mirror the valley in reflectance, explaining the sharp reduction in the intensity of the radiated light in the red region (see also **Fig. 6**).

### BBSs *in vivo*: Skin, Bones, Lymph, and Ova

To evaluate whether the purified BBSs described here *in vitro* can reproduce the hues and saturation observed in *in vivo*, and to test whether the purified BBSs exhibit the same optical properties *in vivo*, we measured the reflectance spectra of live frogs. Reflectance quantifies the spectrally resolved fraction of backscattered light following its interaction with tissue components, including absorption by intra- and extracellular pigments. The frogs’ reflectance spectra show a peak at 520–550 nm and a rapid increase at the far-red/NIR boundary (**Fig. 5**), characteristic of the red-edge in living vegetation^66,67^. Reflectance is markedly lower in the blue portion of the spectrum—likely due to absorption by carotenoids and pteridines^5,9^—and in the red region (660–680 nm) (**Fig. 5**).

Analysis of these reflectance spectra reveals that the reduced reflectance in the red region mirrors the absorbance of pure BBSs, consistently matching their absorbance peaks and spectral features, including a blue-shifted absorbance shoulder (**Fig. 5**). These results suggest that the reflectance spectra of frogs are strongly modulated by the absorbance properties of BBSs.

BBSs generate whole-body green coloration via their distribution throughout the body, including within blood plasma, lymph, and extracellular fluids^4–6^. In many chlorotic frog species, bones exhibit blue-green hues, and in females, oocytes also display green coloration^5,68–81^. To further investigate the distribution of BBSs in these tissues *in vivo*, including within highly pigmented tissues such as blood, we implemented a photoacoustic tomography (PAT) approach. We first verified that the PA spectrum of purified TpBBS matches its absorption spectrum (**Fig. 6A; Supp. Fig. 6**). Next, we imaged an individual *T. pulverata* frog to analyze the spectral properties of internal tissues (**Fig. 6B**). The PA image and spectrum obtained *in vivo* (**Fig. 6A, C**) revealed that TpBBS is present in multiple tissues (673 nm) and can be clearly distinguished from other pigments, such as hemoglobin and melanin at 800 nm (**Fig. 6D, E**). Our results indicate that TpBBS is highly concentrated in bones, lymph, muscles, internal organs, and blood within vessels, reaching values higher than 200 μM in the skin and lymph.

**Fig. 6.**
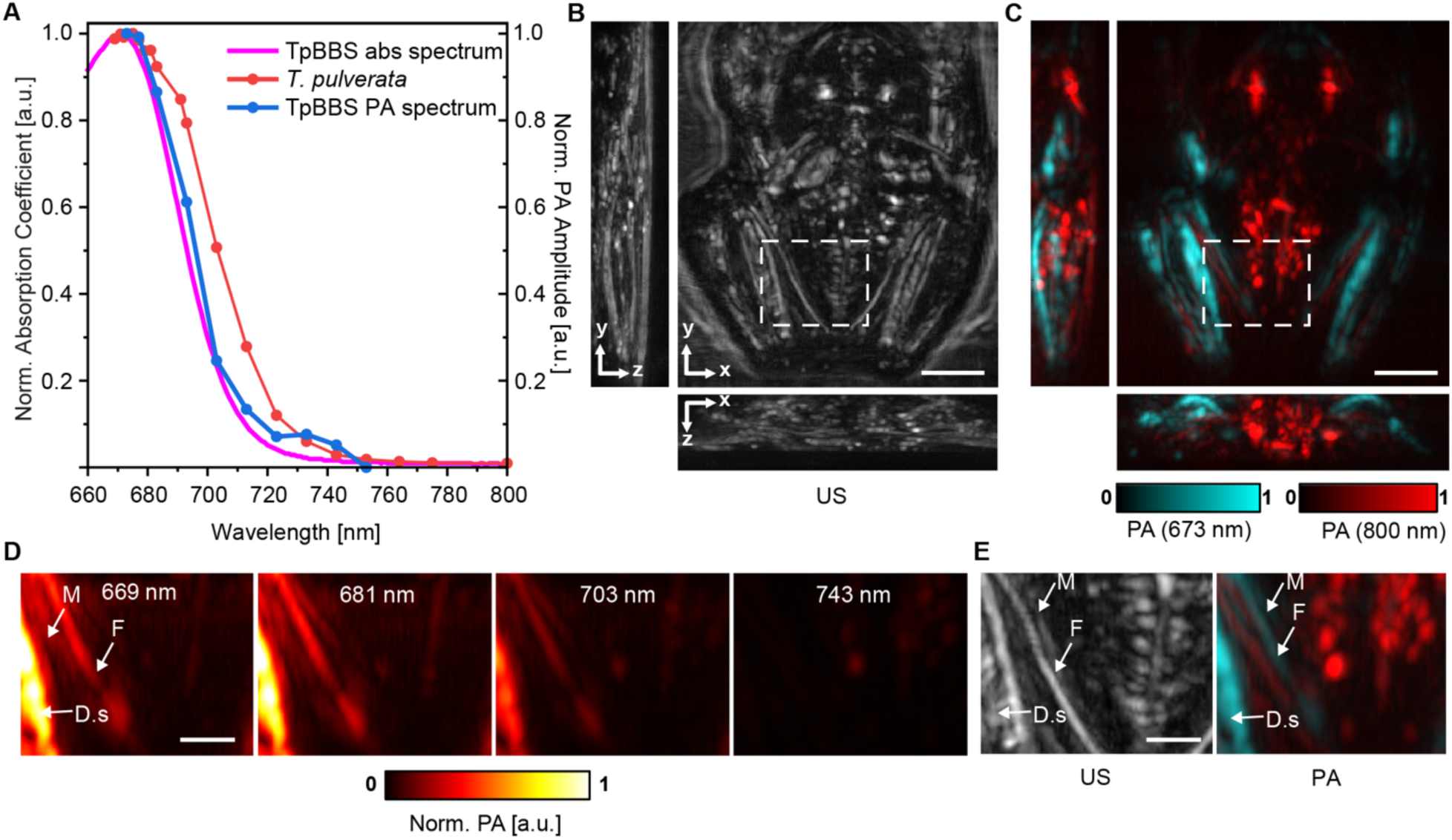
Hyperspectral Photoacoustic Tomography of *T. pulverata* reveals the presence of TpBBS in subcutaneous tissues. **(A)** Average PA amplitude of live frog (blue) and TpBBS-filled tube (light blue/green) compared with the TpBBS’ optical absorption spectrum (red) (See also, **Supp. Fig. 6**). **(B)** Referenced ultrasound (US) image of a sleeping frog, projected along three different dimensions, and **(C)** corresponding overlay of PA images showing TpBBS with 673 nm PA signal (light blue) and other pigments such as hemoglobin and melanin at 800 nm (red). **(D-E)** Zoomed-in region of the US and PA images at increasing wavelengths **(D)**, and US and PA at 673 and 800 nm, respectively **(E)**. TpBBS is distributed across multiple tissues, with high concentrations in the bones (F, femur), lymph, dorsal skin (D.s.), and muscles (M). Because the internal organs of *glassfrogs* are covered by broadband reflectors that backscatter most of the incident light^9,82,83^, the light fluence is attenuated in the dorsal skin immediately overlying the internal organs, which results in weak PA signals. Scale bar: 5 mm in (B–C) and 2 mm in (D–E).

### BBSs produce Near-Infrared fluorescence in frogs

It has been suggested that fluorescence emission may play a role in visual signaling or camouflage in frogs^84–87^. However, many strong fluorophores extracted from amphibian skin have been shown to be quenched and barely detectable *in vivo*^85^. To test whether the fluorescence properties of BBSs are preserved *in vivo*, we imaged multiple individuals using a series of band-pass and long-pass emission filters spanning the first and second near-infrared (SWIR) windows (**Fig. 7A–C; Supp. Fig. 7; Supp. Video 1**).

**Fig. 7.**
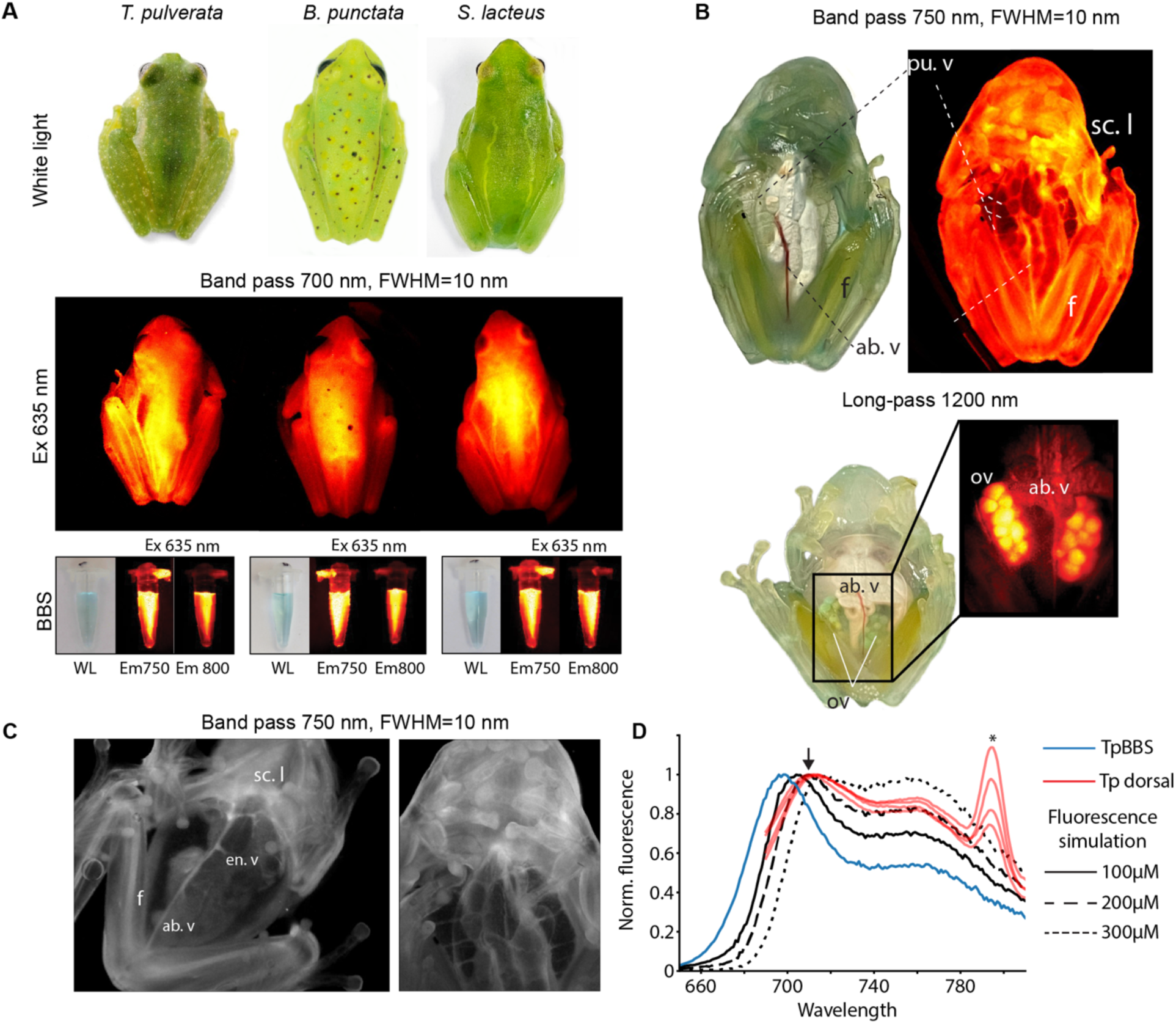
BBSs cause chlorotic hylids and glassfrogs to fluoresce in the NIR when excited in the red/far-red spectral range. Fluorescence emission can be observed in various body regions, including the dorsum **(A)** and the ventral side of the body **(B–C)**, in both sleeping and active frogs (see also **Supp. Video 1**), and in purified BBS samples. Near-infrared images can be visualized either using pseudocolor palettes (A–B) or grayscale (C). BBS are concentrated in subcutaneous lymph, blood plasma, bones, and ova. NIR can be detected from inside the body using band-pass (FWHM=10 nm) and long-pass filters, revealing the distribution of BBS within the organism, and it can be observed at wavelengths as high as 1200 nm. Excitation=635 nm for (A) and 660 nm for (B–C). *ab. v. =* abdominal vein, *en. v. =* enteric vein, *f. =* femur, *ova. =* ovaries, *pu. v.=* pulmonary veins*, sc. l. =* subcutaneous lymph. **(D)** The NIR emission peak is modulated *in vivo* due to fluorescence reabsorption. Monte Carlo simulations of varying BBS concentrations in skin and lymph show that higher concentrations produce a red-shifted emission spectrum compared to pure TpBBS. This trend matches empirical results from multiple individuals, in which the emission peak shifts from 697 nm to approximately 710 nm (arrow). *- Raman peak

In many fluorescent vertebrates, fluorescence is predominantly confined to the UV or visible wavelengths and the distribution of fluorophores is restricted to certain tissues, often within chromatophore cells^3,85,86,88,89^. Contrastingly, NIR emission from chlorotic frogs can be detected broadly across tissues, including the skin, blood, lymph, bones, and ovaries, extending to wavelengths beyond 1200 nm in the short-wavelength infrared (SWIR) window (**Fig. 7C**). Notably, emission is amplified in regions underlain by broadband reflective biological structures, such as the peritoneum and the fasciae surrounding muscles and internal organs. To confirm that the emission *in vivo* follows the same properties as those of isolated BBSs *in vitro*, we obtained fluorescence spectra of adult *T. pulverata.* Emission spectra of live animals are red-shifted compared to those of pure TpBBS (**Fig. 7D**), likely due to light reabsorption, which typically occurs in samples with high concentrations of fluorophores. To assess the effect of BBS concentration on the reabsorption of the emission spectra, we performed a Monte Carlo simulation of the fluorescence emission using a three-layered model that includes an outer skin layer, a middle lymph layer of variable concentrations, and a lower layer composed of a broad-band ideal reflector. Increases in the concentration of BBS in the lymph can effectively modulate the emission, shifting the peak to 710 nm or more, closely matching the spectra obtained *in vivo* (**Fig.7D**). These results confirm that BBSs are responsible for the fluorescence emission of the frogs.

## Discussion

### BBSs and the creation of leaf-like colors

In this work, we characterized the biophysical and optical properties of three independently evolved BBSs from arboreal frogs of the Americas (Hylids and Centrolenids), and confirmed that they can explain the organisms’ leaf-like green coloration, and their fluorescence emission in the NIR window. Our results show that these BBSs are spectrally red-shifted relative to free biliverdin (in the Soret band) and absorb substantially more red light—a feature that generates deep green saturation in a manner analogous to chlorophyll in plants. By increasing the absorption coefficient of the Q band 2.5–3.5-fold and introducing a blue shift in its spectral absorption, chlorotic frogs generate a more saturated green hue than could be achieved with free biliverdin at similar or even higher concentrations.

Frogs that do not rely on biliverdin for coloration instead depend on dermal chromatophore units composed of multiple pigmentary cell types in the skin. Among these, melanin-containing melanophores play a key role by broadly absorbing light and selectively removing red and near-infrared (NIR) wavelengths from backscattered light^1,3^. By contrast, glassfrogs and treefrogs that utilize BBSs, together with a reduction in skin melanin, efficiently subtract red wavelengths while allowing NIR light to be reflected and transmitted with minimal attenuation. This produces an optical phenotype characterized by a sharp increase in reflected light at the transition between the red and NIR regions, known as the red edge, a feature shared with chlorophyll-bearing plants ^66,67^. In the present work, we demonstrated that the spectral features of the Q band of BBSs are detectable in whole animal reflectance spectra. Using hyperspectral PAT, we further showed that the absorption spectral signatures of deep tissues *in vivo*—including bones, ovaries, blood vessels, and lymph—closely match those of BBSs *in vitro*. These results reinforce our conclusion that amphibian coloration arises from a combination of physiological, anatomical, and biochemical mechanisms that integrate seemingly heterogenous systems such as blood metabolism, lymphatic circulation, development, and bone morphogenesis to produce desired optical traits.

The use of biliverdin to generate optical traits in animals requires that the pigment either be stably preserved in tissues for extended periods or produced at rates exceeding its degradation. In non-chlorotic frogs, high doses of biliverdin injected directly into the lymphatic sacs—or artificially elevated biliverdin levels through hemolysis– failed to reproduce the chlorotic phenotype, as the animals rapidly excreted the excess pigment^6^. This observation suggests that elevated biliverdin synthesis alone is insufficient to maintain high concentrations *in vivo* and that additional mechanisms are required for the accumulation of the pigment. In this work, using fluorescence titration experiments performed across multiple concentrations and equilibration times revealed that BBSs bind biliverdin with dissociation constants in the low nanomolar range, and potentially tighter. These affinities are comparable to those of hormone-carrying serpins such as CBG and are 3–4 orders of magnitude higher than those of other vertebrate biliverdin-binding proteins^15,90^. The biliverdin–serpin interaction in BBSs is also 3–5 orders of magnitude stronger than that of an alkaloid-binding serpin from poison dart frogs (K_d_ ∼ 30–50 µM)^91^. Such tight binding likely accounts for the exceptionally high biliverdin concentrations observed in chlorotic frogs. The fact that non-chlorotic frogs cannot accumulate large concentrations of biliverdin might be explained by the lack of an ortholog or any other globulin with sufficient affinity for biliverdin, or a reduced expression of their BBS homologue.

Only one prior study has reported dissociation constants for recombinant BpBBS, estimating a K_d_ of approximately 2.7 μM^15^. This discrepancy likely arises from differences in expression systems (heme oxygenase co-expression), titration conditions (the titration versus binding regime) ^51^, or buffer compositions that could influence binding. Notably, the previously reported absorbance spectrum of recombinant BpBBS differs substantially from that of the endogenous protein^5,15^, exhibiting an 18 nm shift in the Q band and a 4 nm shift in the Soret band, suggesting a different non-native conformation of biliverdin in the binding pocket^7,8,92,93^. In contrast, our current results show that the spectral features of recombinant and endogenous BBSs closely align, suggesting that the biliverdin-binding properties measured *in vitro* here accurately reflect those *in vivo*. These findings also raise the possibility of multiple biliverdin conformations or alternate binding mechanisms.

### Thermal stability and relaxed conformation *in vivo*

We found that BBSs (SlBBS and BpBBS) have melting points above 55 °C, which aligns with the temperatures reported for other serpins in nature^94–98^. In this work, we purified and sequenced the BBS from the glassfrog *T. pulverata* for the first time, and found that TpBBS adopts a hyperstable, relaxed conformation, with a cleaved RCL likely inserted in the serpin fold. The relaxed conformation accounts for the high thermal stability of TpBBS, even at temperatures as high as 95 °C. Currently, there is no clear interpretation or biological role for this increase in stability within the species’ natural history. Furthermore, cleaved serpins are rarely detected *in vivo*, either because they are produced at low levels or are rapidly cleared after inhibiting their target proteases^48,99–102^.

However, their levels can increase in inflammatory cases^103–105^, coagulation disorders^104,106–108^, or tissue injury^109–111^. While the enzyme responsible for cleaving TpBBS *in vivo* is unknown, our sequence alignments suggest that neutrophil elastase may be one of the serine proteases that cleave the reactive center loop. While the biochemical information about amphibian elastases is limited to an ortholog in *Xenopus tropicalis*^112^, its catalytic specificity closely matches that of other vertebrates. Our experimental results showed that human neutrophil elastase cleaves the recombinant TpBBS at M393, consistent with the cleavage site detected in endogenous proteins. This cleavage site is shifted one amino acid toward the C-terminus of the protein relative to the canonical position of the scissile bond position in other serpins^47,52,113^. These results offer an opportunity to investigate structural, biochemical, and evolutionary correlations associated with the length and specificity of the reactive center loop in serpins. It is still unclear whether TpBBS acts as an inhibitor or an alternative protease substrate with purely non-inhibitory functions *in vivo*. Despite undergoing cleavage, TpBBS showed no detectable differences in absorbance between the stressed and relaxed forms, suggesting that serpin cleavage does not alter the ligand binding or the optical effect on the animals. In contrast, corticosteroid- and thyroxine-binding globulins exhibit a marked decrease in hormone affinity following cleavage and transition to the relaxed conformation^59,61,63^. In CBG, this has been interpreted as a regulatory mechanism for corticosteroid release at sites of inflammation, where exposure to target proteases facilitates hormone release^59,103^. By contrast, biliverdin binding in TpBBS appears unaffected by cleavage, suggesting that protease recognition is unlikely to mediate biliverdin release. This stability implies that maintaining biliverdin binding—and the associated spectral tuning—is critical for preserving the coloration and camouflage of *T. pulverata*. A high concentration of cleaved BBSs in normal physiological conditions in a glassfrog offers many avenues to investigate the function and stability of serpins *in vivo*, and to explore the mechanism that permits the retention of optical properties even at high temperatures.

### Other roles of BBSs

By inducing localized skin warming, absorbed radiant energy can be redistributed from superficial layers to deeper tissues through convective transfer via blood circulation^114,115^. In most chlorotic frogs with translucent skin and high concentrations of BBSs in plasma, lymph, muscles, and other deep tissues, BBS may contribute to thermal regulation by absorbing radiant power. Whether frogs can modulate BBS distribution across tissues throughout the day in ways that permit localized heat transfer remains unknown, but further theoretical and experimental work may clarify its potential roles in thermoregulation. Importantly, other roles of BBSs that depend on heterogeneous distribution cannot be ruled out, as uneven distributions of BBSs containing lymph in the lymphatic sacs are common under different physiological conditions.

Alternatively, in this work, we documented that BBSs from treefrogs and the glassfrogs emit fluorescence in the near-infrared portion of the spectrum when excited in the near-UV/violet or the far-red portions of the spectrum. While other amphibian-derived fluorophores in chromatophore cells are likely quenched *in vivo*^85^, BBS fluorescence can be detected from multiple tissues, deep within the organism, such as the vasculature, muscles, ovaries, lymph, and bones. Using fluorescence spectroscopy and Monte Carlo optical modeling, we demonstrated that the emission spectrum of BBSs *in vivo* matches that of the purified BBS which suggests that under physiological conditions, BBSs circulating in the plasma and lymph preserve the same optical traits as when pure.

Whether or not fluorescence emission in the NIR plays a physiological role or is only a byproduct of biliverdin binding to the serpin is unknown. A visual function for NIR emission would require visual systems with photoreceptors sufficiently sensitive to that portion of the spectrum, a condition not documented for purely Opsin-based detectors in nature and one that has been suggested to be physically unlikely^118,119^. However, some organisms, like the deep-sea dragonfish, can use photosensitizers—usually chlorophyll derivatives—to extend their visual sensitivity to the far-red^116^. This phenomenon has also been induced in living rod cells extracted from the axolotl *Ambystoma tigrinum*^117^, and even in mouse models^118^. Currently, there is no information on the visual sensitivity of the frogs studied in this work or evidence suggesting that either the frogs or other interacting species (such as predators, parasites, etc.) might broaden their sensitivity using additional photosensitizers.

A potential visual function in communication requires an increase in contrast with the background that can be perceived by a visual system. Conversely, the leaf-like color produced by BBSs causes a reduction in the contrast, and since the emission of BBSs also overlaps with that of chlorophyll, it is likely that if there is a visual function of fluorescence emission, it would be more related to camouflage. More research is needed to thoroughly evaluate the physiological, ecological, and behavioral effects of fluorescence emission, including other possible non-visual roles.

### Conclusions and opportunities for bioinspiration

BBSs represent a striking example of functional diversification within the serpin superfamily. Their stability, high affinity for biliverdin, and distinctive optical properties generate clear phenotypic effects that can be captured using fast, non-invasive spectroscopic and imaging techniques. Together, these features make the evolution of BBSs a powerful model for investigating novel questions in evolutionary biochemistry and serpin evolution, as well as the biochemical mechanisms that tune the colors of tetrapyrroles. In addition to their evolutionary relevance, the high affinity and stability of BBSs, combined with their fluorescence in the NIR – the optical transparency window– offer opportunities to explore their use as probes for *in vivo* imaging, particularly in contexts where the use of conventional visible-light fluorophores are limited due to the autofluorescence^119–121^. Their strong biliverdin binding can help to circumvent challenges faced by other fluorophores that require endogenous biliverdin in mammals, where biliverdin is rapidly converted to bilirubin^32,122^. Beyond fluorescence, the successful use of PAT to image BBSs *in vivo* demonstrates that their PA spectrum closely matches their *in vitro* absorption spectrum, and that this property is preserved in living systems. This finding suggests that arboreal frog-derived BBSs could be used as a reliable contrast agent for deep-tissue PA imaging, especially for lymphatic applications where current nanoparticle-based optical contrast agents are often considered unstable and toxic^123–128^. Expanding the characterization of BBSs across multiple frog lineages will not only provide new evolutionary insights but also pave the way for harnessing naturally occurring NIR fluorescence for biological imaging.

## Material and methods

### Ethics Declaration

All research procedures and colony care procedures were approved by the Institutional Animal Care and Use Committee of Duke University (protocol numbers A210-18-09 (#78458) and A174-21-08) and California Institute of Technology Protocol IA25-1913.

### Animal species and endogenous BBSs purification

BBSs from adult glassfrogs *Teratohyla pulverata*, and the adult treefrogs *Boana punctata* and *Sphaenorhynchus lacteus*, were extracted and purified as described by Taboada *et al*^5^.

### Size Exclusion Chromatography (SEC)

5 μM of BBS in 100 mM sodium phosphate buffer pH 7.2 were applied to a pre-equilibrated Enrich SEC 650 column (Bio-Rad) at 0.5 ml/min. The protein elution was detected at 280 nm. A SEC calibration curve was prepared with a set of globular protein standards ranging from 1.35 to 670 kDa (Gel Filtration Standard, Bio-Rad).

### Protein Deglycosylation and Crosslinking

Endogenous BBSs were deglycosylated using Reaction Condition Remove-iT PNGase F kit (NEB) with slight modifications from the manufacturer’s instructions. Briefly, 0.5 µL of 10X dithiothreitol (DTT) was added to a 5 µL of BBS solution (20 µg) in 1x phosphate-buffered saline (PBS) pH 7.4 and incubated for 10 minutes at 55 °C to partially denature the protein. 1 µL of 10X Glycobuffer 2 and 2 µL of Remove-iT PNGase F were added, and the samples were incubated at 37 °C overnight. Control conditions with PNGase F-free BBS solutions with 2 µL of HPLC-grade water were incubated at the same temperature conditions to rule out temperature degradation effects. Protein crosslinking was performed using the BS3-d_0_/d_4_ crosslinking agent according to the manufacturer’s protocol.

### *De novo* transcriptome assembly

One specimen of *T. pulverata* was euthanized, and the liver was immediately excised and preserved in 1.5 mL RNA*later* (Thermo Scientific). Tissue was homogenized using Zirconium beads, and total RNA was extracted using the *Quick*-RNA Miniprep Kit (Zymo Research) following the manufacturer’s instructions. The extracted RNA was sent to Novogene for sequencing, using 150-bp pair-end reads. The raw sequences were cleaned using Trimmomatic v0.39^129^ run in pair-end mode (3’ end quality = 20; sliding window mean quality = 20, and minimum length = 36). Cleaned sequences were assembled using Trinity v2.15.2^130^.

### Mass Spectrometry and Protein Sequencing

BBSs were subjected to SDS-PAGE and stained with Coomassie brilliant blue (CBB R250). The CBB-stained gel bands were cut out, destained, reduced with DTT, and alkylated with iodoacetamide, followed by digestion with Trypsin Gold (Promega). The trypsinized peptides were lyophilized and reconstituted in 20 μL of 60% acetonitrile/0.1% TFA. 2 μL of each sample was analyzed using a 30 min gradient by nanoLC-MS/MS on an Orbitrap Fusion Lumos (Thermo Scientific). MS/MS was acquired in a high-resolution, data-dependent mode on an Orbitrap. Mass spectrometry data were analyzed using the software PEAKS^131^. The identified peptides were matched against the protein database generated from the *de novo* assembly of the transcriptome of *T. pulverata.* To identify the site of cleavage of the C-terminus region of the endogenous TpBBS, native TpBBS was analyzed in LTQ Orbitrap XL Hybrid Ion Trap-Orbitrap Mass Spectrometer (Thermo Scientific) paired with a Waters Acquity UPLC. MS data obtained from the Orbitrap were manually analyzed.

### MALDI-TOF

Molecular weight of purified BBSs was determined by MALDI-TOF on an Autoflex TOF/TOF mass spectrometer (Bruker). The MALDI matrix was prepared by combining 50 mg sinapinic acid (SA), 1 ml HPLC grade water, 1 ml acetonitrile, with 0.1% TFA. Samples were prepared mixing 1 µl of protein with 1 µl of matrix, and co-crystalized at room temperature. For the molecular mass determination, a mixture consisting of four standard proteins- insulin, ubiquitin I, cytochrome C, myoglobin (Protein Calibration Standard I, Bruker)- ranging from 4 to 20 kDa was used.

### Sequence alignment and structure prediction

All protein sequences were aligned using T-Coffee^132^. The resulting multiple sequence alignment was manually refined using Jalview^133^. For estimating sequence identity and similarity, the N-terminal region of the serpins was not considered. The sequence alignment was visualized with Sequence Manipulation Suite (Color Align Properties)^134^. 3D structure of the BBS was predicted using AlphaFold3^135^.

### Spectroscopy measurements

The absorbance spectrum of the protein and ligand was collected from 250 nm to 850 nm in Jasco V-750 UV-Vis spectrophotometer. The final spectrum is background corrected with the appropriate buffer. BBS (1-2 µM) emission fluorescence was collected by exciting the samples at 390 nm or 630 nm and monitoring the emission from 650 nm to 750 nm in Edinburgh FS5 fluorimeter, with excitation and emission bandwidths of 5 nm. The presented fluorescence spectrum is averaged over three accumulations.

The relative quantum yield of BBSs was estimated as described in literature^10,136^. The fluorescence emission of BBSs and Cy5.5 (absorbance at 650 nm ranging from 0.01-0.1) was measured from 670-900 nm by exciting the samples at 650 nm. Cy5.5 (QY = 0.28) dissolved in 1X PBS was used as the standard, as its excitation and emission spectra overlap with those of BBSs^5,15^.

### Thermal unfolding

When the BBSs are unfolded, the release of biliverdin is accompanied by a shift in the Soret band of biliverdin from 390 nm to 376 nm. Thermal unfolding of BBSs was monitored by measuring the absorbance at 376 nm and 390 nm BBS in 1X PBS (A_390_ = 0.16) was gradually heated from 20 °C to 90 °C at a rate of 1 °C/min, and the absorbance changes (A390/A376) were monitored in an Agilent Cary UV-Vis spectrophotometer or an Edinburgh FS5 spectrometer connected to the Peltier controller. The melting point was calculated from the first derivative of the absorbance signal as a function of time, and the peak of the derivative corresponds to the melting temperature of the protein.

### Cloning, protein overexpression, and purification

The coding sequence corresponding to the TpBBS, BpBBS, and SlBBS were synthesized from GeneArt (Germany). BBS genes were subcloned into pBG106 expression vector (Center for Structural Biology, Vanderbilt University) through In-fusion HD Cloning (Takara Bio Inc.). TpBBS and SlBBS were overexpressed in *E. coli* Rosetta2(DE3) cells. The transformed bacterial cells were cultured at 37 °C to an OD_600_ of 0.6, then the overexpression was induced with 0.2 mM IPTG for 16 h at 22 °C with constant shaking. BpBBS was overexpressed in *E. coli* Rosetta2 (DE3) cells grown to OD_600_ = 1.0, then induced with 1 mM IPTG for 16 h at 16 °C with constant shaking. Cells were pelleted and resuspended to homogeneity in lysis buffer (20 mM Tris pH 8, 150 mM NaCl, 2 protease inhibitor tablets (Roche)). The cells were lysed by sonication, and the cell lysate was clarified by centrifuging at 10,000 xg for 40 min. The lysate was loaded onto the Nickel Sepharose 6 Fast Flow affinity column (Cytiva) pre-equilibrated with the lysis buffer. For SlBBS and TpBBS, the non-specific impurities were washed with lysis buffer containing 50 mM imidazole, and the bound protein was eluted with lysis buffer containing 300 mM imidazole. The eluted protein was dialyzed against QA buffer (20 mM Tris pH 8, 25 mM NaCl, 1 mM DTT) overnight at 4 °C. The dialyzed protein was loaded onto an Uno Q6 (Bio-Rad) column connected to Bio-Rad NGC FPLC and eluted with a salt gradient ranging from 25-500 mM NaCl. The peak fractions containing BBS were pooled together, dialyzed against storage buffer (20 mM Tris pH 8, 50 mM NaCl, 10% glycerol) overnight. For BpBBS, non-specific impurities were washed with lysis buffer containing 20 mM imidazole and eluted with 300 mM imidazole. The eluted protein was buffer-exchanged into 1X PBS. The protein was then concentrated, aliquoted into small fractions and flash frozen.

### Relaxed TpBBS preparation

The endogenous TpBBS was identified in the relaxed confirmation with an elastase recognition site in the reactive center loop. The relaxed TpBBS was prepared by incubating the recombinant TpBBS with human neutrophil elastase (Innovative Research, Inc.) at 1:3 ratio at 25 °C for 6 h in 1X PBS. Protein cleavage was confirmed by western blotting. Anti-his antibody (Ab18184, Abcam Ltd.) was used to confirm the proteolytic cleavage.

### Determination of biliverdin, apo-BBSs, and holo-BBSs concentrations

Apo-BBSs concentrations were determined by measuring absorbance at 280 nm^137^ (ε_280, TpBBS_ = 33,350 M^-1^ cm^-1^, ε_280, BpBBS_ = 28,880 M^-1^ cm^-1^, ε_280, SlBBS_ = 35,870 M^-1^ cm^-1^). For the quantification of free biliverdin (Cayman Chemical) in 1X PBS, we used the extinction coefficient at the Soret band (c_376 nm_ = 39,900 M^-1^cm^-1^) and the Q band (c_672 nm_ of 11,200 M^-1^cm^-1^) as determined in this study (**Supp. Fig. 3**). A known mass (10 mg, 95% purity) of biliverdin was dissolved in DMSO at 9.5 mg/mL, and further diluted to different concentrations in 1X PBS, and the absorbance spectrum was recorded from 250 to 850 nm. The molar extinction coefficient of biliverdin in 1X PBS was determined from the slope of the absorbance vs concentration plot (**Supp Fig. 3**). The extinction coefficients of biliverdin bound to the BBSs at the Soret and Q bands were determined using recombinant proteins, wherein a fixed concentration of biliverdin was titrated with increasing concentrations of apo-BBS until no further changes in absorption were observed.

### Ligand binding and determination of dissociation constants (Kd)

Dilutions of recombinant BBS were prepared in a flat black 96 well plate (Corning) spanning 0.2–30 nM in 1X PBS, to capture the K_d_ in the binding regime^51^. 15 nM biliverdin was added to each well, and fluorescence emission was collected at 697 nm (rTpBBS), 710 nm (rBpBBS), and 707 nm (rSlBBS) with 20 nm bandwidths for excitation and emission wavelength at an optimal gain. Fluorescence intensities were measured using a Tecan Spark plate reader with a small humidity cassette filled with ultrapure water to reduce evaporative sample loss, and scans were taken once every hour for 24 h at room temperature. The equilibrium dose-response curve was plotted (normalized for relative fluorescence intensity vs. concentration) and the K_d_ value was determined by fitting the data with the Sigmoidal curve equation on GraphPad Prism version 10.6.1 for Windows, GraphPad Software, Boston, Massachusetts USA, www.graphpad.com.

### *In vivo* spectroscopy

We measured the reflectance spectra of the frogs following the methodology reported by Taboada *et al*.^9^. Light from the dorsum of the frog was measured using an integrating sphere (Ocean Optics ISP-REF, port diameter 10.32 mm) connected to a multichannel spectroradiometer (Ocean Optics, USB 2000). We used the built-in tungsten halogen lamp within the Integrating sphere ISP-REF for the light source. To obtain reflectance measurements, a Spectralon diffuse 100% reflectance standard was used (Labsphere Inc.) to calibrate the device before each spectrum acquisition.

The *in vivo* fluorescence emission was measured in adult specimens of *Teratohyla pulverata.* To simulate the effect of BBSs absorption on the frog’s emission spectra, we performed a 2D Monte Carlo simulation using a three-layered system consisting of a broad-band reflector, a subcutaneous lymph layer with varying BBS concentrations, and an outer skin layer. A single fluorescent point source was defined for the simulation in the BBS-filled medium. The simulation was performed using the MCXLAB simulation package, which is the native MEX version of MCX for MATLAB^138,139^. The simulation grid comprised a 300×300 matrix of isotropic pixels with a pixel width of ∼3.3 µm, corresponding to a total grid area of 1 mm^2^. The medium was split into three layers; an open-air background layer, a BBS-containing layer (split into skin and lymph), and an ideal specular reflector layer. The BBS skin and lymph layers were each 0.1 mm thick, with the reflector layer below it also being 0.1 mm thick. Four optical properties were defined for each layer in the model: absorption coefficient, scattering coefficient, scattering anisotropy (g), and refractive index. The scattering and absorption coefficients for the BBS containing layers were based on the empirically measured absorption and scattering spectra of BBS. The scattering anisotropy and refractive index were defined as 0.9 and 1.4 for each layer in the model, respectively. 171 simulations were performed independently and subsequently, to model emission using the optical properties of BBS from 650 nm to 820 nm in increments of 1 nm. For each simulation, a single fluorescent-point-source pulse emitted light from the center of the BBS-filled layer, and the cumulative light reaching a point detector defined in the open-air layer was calculated. The intensity of the initial point source of light was defined using the empirical emission spectrum of BBS. 50 million photon packets were modelled for each simulation, with a temporal resolution of 0.1 ps for a duration of 5 ps. Following the 171 simulations, the resulting relative emission profile was plotted based on the total light reaching the detector for each of the wavelengths simulated.

To estimate the skin’s optical properties, we applied an inverse adding–doubling (IAD) approach to reflectance and transmittance spectra acquired using an integrating sphere with a 10-mm port and a 1-mm incident light beam on the skin surface^140^. In all the simulations, we assumed that the asymmetry parameter is 0.9 for the skin.

### NIR fluorescence imaging

Fluorescent images were acquired using an NIR high-sensitivity InGaAs camera with a spectral range of 600 nm to 1.7 μm (Ninox 640 II, Raptor Photonics). Excitation was provided by an LED source with a 660 nm peak wavelength and a 20 nm full-width-at-half-maximum (FWHM) (Thorlabs, M660L4). Emission was detected using band-pass filters (700 ± 10 nm, 750 ± 10 nm, 800 ± 10 nm) and long-pass filters (1200 nm cut-on).

### Three-dimensional photoacoustic (PA) and ultrasound (US) imaging

For multispectral wide-field imaging of the frogs, we employed a custom-built linear-array-based 3D PAT system based on diffractive acoustic tomography^141^. The system allowed for isotropic three-dimensional PA and US imaging of whole frogs. The system used a linear-array transducer (L7-4, Philips) combined with an optically transparent single-slit to enhance elevational resolution. Acoustic data from the transducer were acquired by a 256-channel data acquisition (DAQ) system (Vantage 256, Verasonics, Inc.). For PA imaging, an Nd:YAG laser generated pulses (10 ns pulse width, and 10 Hz pulse repetition rate (PRR)) that were delivered to the frog through a dual-branch line fiber bundle (Dolan Jenner) from the ventral side. Before coupling into the line fiber bundle, a beam sampler directed <5 % of the laser pulse energy into a power meter (Ophir) later used for energy normalization. The Nd:YAG laser pumped an optical parametric oscillator (OPO) (Continuum, Inc.) to allow for tunable wavelength control from 680-1000 nm. Laser pulsing and ultrasonic detection are controlled by an embedded device with FPGA module (myRIO-1900, NI). Each volumetric dataset for each wavelength used to image the frogs required ∼30 seconds, with 300 scanning steps (step size, 0.1 mm) defining a volume of 4 × 3 × 2 cm, covering the entire frog, with an isotropic spatial resolution of approximately 400 µm. To evaluate if BBS absorption *in vivo* matches the spectral distribution of purified BBS, we injected purified TpBBS into optically and acoustically transparent micro dialysis tubes and obtained the PA spectra for comparison.

## Acknowledgments

This work was partially sponsored by the David and Lucile Packard Fellowship for Science and Engineering (CT), Beckman Young Investigator Award (CT), start-up funds from Vanderbilt University and the California Institute of Technology (CT), National Geographic Society grant NGS-65348R-19 (JD, CT), the United States National Institutes of Health (NIH) grants RF1 NS115581, R01 NS111039, R01 EB028143, R01 DK139109, R01 DK052985 (JY), The United States National Science Foundation (NSF) CAREER award 2144788 (JY), Chan Zuckerberg Initiative Grant 2024-349531 (JY), American Heart Association Collaborative Science Award 25CSA1417550 (JY).

## Author contributions

Conceptualization: C.T., R.V., G.H. Methodology: C.T., R.V., G.H., T.V., L.M. Investigation: R.V., T.V., G.H., L.M., J.D., W.W., P.L., E.G., C.T. Data Analysis: R.V., T.V., G.H., L.M., C.T. Writing – original draft: C.T., R.V., and G.H. Writing – review and editing: C.T., R.V., T.V., G.H., L.M., J.D., W.W., P.L., E.G., S.J., J.Y., C.T. Supervision: C.T. Funding Acquisition: C.T., J.D., J.Y.

## Data Availability

Data supporting the findings of this study are available within the article and its supplementary materials.

## References

1. Bagnara, J. T. & Matsumoto, J. Comparative Anatomy and Physiology of Pigment Cells in Nonmammalian Tissues. in The Pigmentary System 11–59 (John Wiley & Sons, Ltd, 2006).

2. Taylor, J. D. & Bagnara, J. T. Dermal Chromatophoies. Am. Zool. 12, 43–62 (1972).

3. Bagnara, J. T., Taylor, J. D. & Hadley, M. E. The Dermal Chromatophore Unit. J. Cell Biol. 38, 67–79 (1968).

4. Barrio, A. Cloricia fisiológica en batracios anuros. Physis 25, 137–142 (1965).

5. Taboada, C. et al. Multiple origins of green coloration in frogs mediated by a novel biliverdin-binding serpin. Proc. Natl. Acad. Sci. 117, 18574–18581 (2020).

6. Jones, D. A. Green Pigmentation in Neotropical Frogs. PhD Thesis. (The University of Florida, 1967).

7. Falk, H. The Chemistry of Linear Oligopyrroles and Bile Pigments. (Springer Science & Business Media, 2012).

8. Taniguchi, M. & Lindsey, J. S. Absorption and fluorescence spectra of open-chain tetrapyrrole pigments–bilirubins, biliverdins, phycobilins, and synthetic analogues. J. Photochem. Photobiol. C Photochem. Rev. 55, 100585 (2023).

9. Taboada, C. et al. Glassfrogs conceal blood in their liver to maintain transparency. Science 378, 1315–1320 (2022).

10. Rodriguez, E. A. et al. A far-red fluorescent protein evolved from a cyanobacterial phycobiliprotein. Nat. Methods 13, 763–769 (2016).

11. Ghosh, S. et al. Blue protein with red fluorescence. Proc. Natl. Acad. Sci. 113, 11513–11518 (2016).

12. Fischer, A. J. & Lagarias, J. C. Harnessing phytochrome’s glowing potential. Proc. Natl. Acad. Sci. 101, 17334–17339 (2004).

13. Shu, X. et al. Mammalian Expression of Infrared Fluorescent Proteins Engineered from a Bacterial Phytochrome. Science 324, 804–807 (2009).

14. Shcherbakova, D. M. & Verkhusha, V. V. Near-infrared fluorescent proteins for multicolor in vivo imaging. Nat. Methods 10, 751–754 (2013).

15. Manoilov, K. Yu., Ghosh, A., Almo, S. C. & Verkhusha, V. V. Structural and Functional Characterization of a Biliverdin-Binding Near-Infrared Fluorescent Protein From the Serpin Superfamily. J. Mol. Biol. 434, 167359 (2022).

16. Golovynskyi, S. et al. Optical windows for head tissues in near-infrared and short-wave infrared regions: Approaching transcranial light applications. J. Biophotonics 11, e201800141 (2018).

17. Jacques, S. L. Origins of Tissue Optical Properties in the UVA, Visible, and NIR Regions. in Advances in Optical Imaging and Photon Migration OPC364 (Optica Publishing Group, 1996).

18. Jacques, S. L. Optical properties of biological tissues: a review. Phys. Med. Biol. 58, R37 (2013).

19. Krause, G. H. & Weis, E. Chlorophyll fluorescence as a tool in plant physiology. Photosynth. Res. 5, 139–157 (1984).

20. Krause, G. H. & Weis, E. Chlorophyll Fluorescence and Photosynthesis: The Basics. Annu. Rev. Plant Biol. 42, 313–349 (1991).

21. Oliinyk, O. S., Chernov, K. G. & Verkhusha, V. V. Bacterial Phytochromes, Cyanobacteriochromes and Allophycocyanins as a Source of Near-Infrared Fluorescent Probes. Int. J. Mol. Sci. 18, 1691 (2017).

22. Crawford, J. M., Ransil, B. J., Potter, C. S., Westmoreland, S. V. & Gollan, J. L. Hepatic disposition and biliary excretion of bilirubin and bilirubin glucuronides in intact rats. Differential processing of pigments derived from intra- and extrahepatic sources. J. Clin. Invest. 79, 1172–1180 (1987).

23. McDonagh, A. F., Lightner, D. A., Kar, A. K. & Norona, W. S. Hepatobiliary excretion of biliverdin isomers and C10-substituted biliverdins in Mrp2-deficient (TR−) rats. Biochem. Biophys. Res. Commun. 293, 1077–1083 (2002).

24. Cornelius, C. E. Biliverdin in Biological Systems. in Biliverdin in Biological Systems (eds Ryder, O. A. & Byrd, M. L.) 321–334 (Springer, Berlin, Heidelberg, 1984).

25. Hirota, K. Urinary excretion of alpha- and beta-isomers of biliverdin-IX in humans. Biol. Pharm. Bull. 18, 481–484 (1995).

26. Cornelius, C. E. & Bruss, M. L. Hepatic bile pigment excretion and erythrocyte turnover in various species. Vet. Clin. Pathol. 9, 15–20 (1980).

27. Hirota, K., Yamamoto, S. & Itano, H. A. Urinary excretion of isomers of biliverdin after destruction in vivo of haemoproteins and haemin. Biochem. J. 229, 477–483 (1985).

28. Barrowman, J. A., Bonnett, R. & Bray, P. J. Metabolism of biliverdin. Biliary excretion of bile pigments after intravenous injection of biliverdin isomers. Biochim. Biophys. Acta 444, 333–337 (1976).

29. Ramonas, L. M., McDonagh, A. F. & Palma, L. A. A flow-cell method for measuring hepatic excretion of pigments continuously in rats. J. Pharmacol. Methods 5, 149–164 (1981).

30. Bagnara, J. T., Fernandez, P. J. & Fujii, R. On the blue coloration of vertebrates. Pigment Cell Res. 20, 14–26 (2007).

31. Kratochwil, C. F. & Mallarino, R. Mechanisms Underlying the Formation and Evolution of Vertebrate Color Patterns. Annu. Rev. Genet. 57, 135–156 (2023).

32. Kasatkina, L. A. et al. Deep-tissue high-sensitivity multimodal imaging and optogenetic manipulation enabled by biliverdin reductase knockout. Nat. Commun. 16, 6469 (2025).

33. Shiels, R. G. et al. Pharmacokinetics of bilirubin-10-sulfonate and biliverdin in the rat. Eur. J. Pharm. Sci. Off. J. Eur. Fed. Pharm. Sci. 159, 105684 (2021).

34. van Dijk, R. et al. Biliverdin Reductase inhibitors did not improve severe unconjugated hyperbilirubinemia in vivo. Sci. Rep. 7, 1646 (2017).

35. Maiti, A. et al. Structural and photophysical characterization of the small ultra-red fluorescent protein. Nat. Commun. 14, 4155 (2023).

36. Kobachi, K. et al. Biliverdin Reductase-A Deficiency Brighten and Sensitize Biliverdin-binding Chromoproteins. Cell Struct. Funct. 45, 131–141 (2020).

37. Montecinos-Franjola, F., Lin, J. Y. & Rodriguez, E. A. Fluorescent proteins for in vivo imaging, where’s the biliverdin? Biochem. Soc. Trans. 48, 2657–2667 (2020).

38. Peng, H.-P., Lee, K. H., Jian, J.-W. & Yang, A.-S. Origins of specificity and affinity in antibody–protein interactions. Proc. Natl. Acad. Sci. 111, E2656–E2665 (2014).

39. Puett, D. & Angelova, K. Determining the affinity of hormone-receptor interaction. Methods Mol. Biol. 590, 1–20 (2009).

40. Esparza, T. J., Martin, N. P., Anderson, G. P., Goldman, E. R. & Brody, D. L. High affinity nanobodies block SARS-CoV-2 spike receptor binding domain interaction with human angiotensin converting enzyme. Sci. Rep. 10, 22370 (2020).

41. Schellenberg, M. J., Petrovich, R. M., Malone, C. C. & Williams, R. S. Selectable high-yield recombinant protein production in human cells using a GFP/YFP nanobody affinity support. Protein Sci. 27, 1083–1092 (2018).

42. Unson, C. G., Wu, C.-R., Sakmar, T. P. & Merrifield, R. B. Selective Stabilization of the High Affinity Binding Conformation of Glucagon Receptor by the Long Splice Variant of Gαs*. J. Biol. Chem. 275, 21631–21638 (2000).

43. Zhang, J.-L., Simeonowa, I., Wang, Y. & Sebald, W. The high-affinity interaction of human IL-4 and the receptor alpha chain is constituted by two independent binding clusters. J. Mol. Biol. 315, 399–407 (2002).

44. Esbenshade, T. A. et al. The histamine H3 receptor: an attractive target for the treatment of cognitive disorders. Br. J. Pharmacol. 154, 1166–1181 (2008).

45. Rajapaksha, H. & Forbes, B. E. Ligand-Binding Affinity at the Insulin Receptor Isoform-A and Subsequent IR-A Tyrosine Phosphorylation Kinetics are Important Determinants of Mitogenic Biological Outcomes. Front. Endocrinol. 6, 107 (2015).

46. Basu, S. et al. Determination of Binding Affinity of Antibodies to HIV-1 Recombinant Envelope Glycoproteins, Pseudoviruses, Infectious Molecular Clones, and Cell-Expressed Trimeric gp160 Using Microscale Thermophoresis. Cells 13, 33 (2023).

47. Marijanovic, E. M. et al. Reactive centre loop dynamics and serpin specificity. Sci. Rep. 9, 3870 (2019).

48. Gettins, P. G. W. Serpin Structure, Mechanism, and Function. Chem. Rev. 102, 4751–4804 (2002).

49. Huntington, J. A. Serpin structure, function and dysfunction. J. Thromb. Haemost. 9, 26–34 (2011).

50. Fulton, K. F. et al. The High Resolution Crystal Structure of a Native Thermostable Serpin Reveals the Complex Mechanism Underpinning the Stressed to Relaxed Transition*. J. Biol. Chem. 280, 8435–8442 (2005).

51. Jarmoskaite, I., AlSadhan, I., Vaidyanathan, P. P. & Herschlag, D. How to measure and evaluate binding affinities. eLife 9, e57264 (2020).

52. Sanrattana, W., Maas, C. & de Maat, S. SERPINs—From Trap to Treatment. Front. Med. 6, (2019).

53. Kumar, A. Bayesian phylogeny analysis of vertebrate serpins illustrates evolutionary conservation of the intron and indels based six groups classification system from lampreys for ∼500 MY. PeerJ 3, e1026 (2015).

54. Silverman, G. A. et al. The Serpins Are an Expanding Superfamily of Structurally Similar but Functionally Diverse Proteins. J. Biol. Chem. 276, 33293–33296 (2001).

55. Carrell, R. W., Pemberton, P. A. & Boswell, D. R. The serpins: evolution and adaptation in a family of protease inhibitors. Cold Spring Harb. Symp. Quant. Biol. 52, 527–535 (1987).

56. Whisstock, J. C. et al. Serpins Flex Their Muscle: II. Structural insights into target peptidase recognition, polymerization, and transport functions*. J. Biol. Chem. 285, 24307–24312 (2010).

57. Law, R. H. P. et al. An overview of the serpin superfamily. Genome Biol. 7, 216 (2006).

58. Whisstock, J. C. & Bird, P. I. Serpin Structure and Evolution. vol. 501 (Academic PRess, 2011).

59. Chan, W. L., Carrell, R. W., Zhou, A. & Read, R. J. How changes in affinity of corticosteroid-binding globulin modulate free cortisol concentration. J. Clin. Endocrinol. Metab. 98, 3315–3322 (2013).

60. Carrell, R., Qi, X. & Zhou, A. Serpins as hormone carriers: modulation of release. Methods Enzymol. 501, 89–103 (2011).

61. Zhou, A., Wei, Z., Read, R. J. & Carrell, R. W. Structural mechanism for the carriage and release of thyroxine in the blood. Proc. Natl. Acad. Sci. U. S. A. 103, 13321–13326 (2006).

62. Zhou, A. et al. A redox switch in angiotensinogen modulates angiotensin release. Nature 468, 108–111 (2010).

63. Qi, X. et al. Allosteric Modulation of Hormone Release from Thyroxine and Corticosteroid-binding Globulins. J. Biol. Chem. 286, 16163–16173 (2011).

64. Meyer, J. F., Bieth, J. & Metais, P. On the inhibition of elastase by serum. Some distinguishing properties of alpha1-antitrypsin and alpha2-macroglobulin. Clin. Chim. Acta Int. J. Clin. Chem. 62, 43–53 (1975).

65. Cohen, A. B. The interaction of alpha-1-antitrypsin with chymotrypsin, trypsin and elastase. Biochim. Biophys. Acta 391, 193–200 (1975).

66. Horler, D. N. H., Dockray, M. & Barber, J. The red edge of plant leaf reflectance. Int. J. Remote Sens. 4, 273–288 (1983).

67. Blount, C. C. Near infrared reflectance in Anura. PhD Thesis (The University of Manchester, Manchester, UK, 2018).

68. Guayasamin, J. M. et al. Phylogenetic systematics of Glassfrogs (Amphibia: Centrolenidae) and their sister taxon Allophryne ruthveni. Zootaxa 2100, (2009).

69. Twomey, E., Delia, J. & Castroviejo-Fisher, S. A review of Northern Peruvian glassfrogs (Centrolenidae), with the description of four new remarkable species. Zootaxa 3851, (2014).

70. Catenazzi, A., May, R. V., Lehr, E., Gagliardi-Urrutia, G. & Guayasamin, J. M. A new, high-elevation glassfrog (Anura: Centrolenidae) from Manu National Park, southern Peru. Zootaxa 3388, 56–68 (2012).

71. Castroviejo-Fisher, S., Vilà, C., Ayarzagüena, J., Blanc, M. & Ernst, R. Species diversity of Hyalinobatrachium glassfrogs (Amphibia: Centrolenidae) from the Guiana Shield, with the description of two new species. Zootaxa 3132, 1–55 (2011).

72. Berneck, B. V. M., Haddad, C. F. B., Lyra, M. L., Cruz, C. A. G. & Faivovich, J. The Green Clade grows: A phylogenetic analysis of Apalstodiscus (Anura; Hylidae). Mol. Phylogenet. Evol. 97, 213–223 (2016).

73. Garda, A. A., Santana, D. J. & São-Pedro, V. D. A. Taxonomic characterization of Paradoxical frogs (Anura, Hylidae, Pseudae): geographic distribution, external morphology, and morphometry. Zootaxa 2666, 1–28 (2010).

74. Hoogmoed, M. S. Notes on the herpetofauna of Surinam : VI. Resurrection of Hyla ornatissima Noble (Amphibia, Hylidae) and remarks on related species of green tree frogs from the Guiana area. Zool. Verh. 172, 3–46 (1979).

75. Channing, A. et al. Taxonomy of the super-cryptic Hyperolius nasutus group of long reed frogs of Africa (Anura: Hyperoliidae), with descriptions of six new species. Zootaxa 3620, 301–350 (2013).

76. Preez, L. H. D. & Carruthers, V. Frogs of Southern Africa: A Complete Guide. (Struik Nature, 2017).

77. Glaw, F. & Vences, M. A Field Guide to the Amphibians and Reptiles of Madagascar. (2007).

78. Blommers-Schlösser, R. M. A. Biosystematics of the Malagasy Frogs. II. The Genus Boophis (Rhacophoridae). Bijdr. Tot. Dierkd. 49, 261-p4 (1979).

79. Biju, S. D. et al. Taxonomic review of the tree frog genus Rhacophorus from the Western Ghats, India (Anura: Rhacophoridae), with description of ontogenetic colour changes and reproductive behavior. Zootaxa 3636, 257–289 (2013).

80. Harvey, M. B., Pemberton, A. J. & Smith, E. N. New and poorly known parachuting frogs (Rhacophoridae : Rhacophorus) from Sumatra and Java. Herpetol. Monogr. 16, 46–92 (2002).

81. Onn, C. K. A Field Guide to the Frogs of Borneo. Third edition. Copeia 106, 396–397 (2018).

82. Cisneros-Heredia, D. F. & McDiarmid, R. W. Revision of the characters of Centrolenidae (Amphibia: Anura: Athesphatanura), with comments on its taxonomy and the description of new taxa of glassfrogs. Zootaxa 1572, 1–82 (2007).

83. Rong, Q. et al. Label-free photoacoustic imaging of glassfrog development. Photoacoustics 46, 100773 (2025).

84. Taboada, C. et al. Naturally occurring fluorescence in frogs. Proc. Natl. Acad. Sci. 114, 3672–3677 (2017).

85. Taboada, C., Brunetti, A. E., Alexandre, C., Lagorio, M. G. & Faivovich, J. Fluorescent Frogs: A Herpetological Perspective. *South Am*. J. Herpetol. 12, 1–13 (2017).

86. Friedrich, S., Schwager, M., Heß, M., Glaw, F. & Lehmann, T. Evidence for fluorescence-supported species recognition in syntopic harvestmen. Sci. Rep. 16, 2631 (2026).

87. Whitcher, C. et al. Evidence for ecological tuning of anuran biofluorescent signals. Nat. Commun. 15, 8884 (2024).

88. Wucherer, M. F. & Michiels, N. K. Regulation of red fluorescent light emission in a cryptic marine fish. Front. Zool. 11, 1 (2014).

89. Prötzel, D., Heß, M., Schwager, M., Glaw, F. & Scherz, M. D. Neon-green fluorescence in the desert gecko Pachydactylus rangei caused by iridophores. Sci. Rep. 11, 297 (2021).

90. Ghosh, S. et al. Modulation of biliverdin dynamics and spectral properties by Sandercyanin. RSC Adv. 12, 20296–20304.

91. Alvarez-Buylla, A. et al. Binding and sequestration of poison frog alkaloids by a plasma globulin. eLife 12, e85096 (2023).

92. Iturraspe, J. B., Bari, S. & Frydman, B. Total synthesis of ‘extended’ biliverdins. The relation between their conformation and their spectroscopic properties. J. Am. Chem. Soc. 111, 1525–1527 (1989).

93. Braslavsky, S. E., Holzwarth, A. R. & Schaffner, K. Solution Conformations, Photophysics, and Photochemistry of Bile Pigments; Bilirubin and Biliverdin, Dimethyl Esters and Related Linear Tetrapyrroles. Angew. Chem. Int. Ed. Engl. 22, 656–674 (1983).

94. Benning, L. N., Whisstock, J. C., Sun, J., Bird, P. I. & Bottomley, S. P. The human serpin proteinase inhibitor-9 self-associates at physiological temperatures. Protein Sci. Publ. Protein Soc. 13, 1859–1864 (2004).

95. Huntington, J. A. & Carrell, R. W. The Serpins: Nature’s Molecular Mousetraps. Sci. Prog. 84, 125–136 (2001).

96. Irving, J. A. et al. The 1.5 Å Crystal Structure of a Prokaryote Serpin: Controlling Conformational Change in a Heated Environment. Structure 11, 387–397 (2003).

97. Dafforn, T. R., Mahadeva, R., Elliott, P. R., Sivasothy, P. & Lomas, D. A. A Kinetic Mechanism for the Polymerization of α1-Antitrypsin. J. Biol. Chem. 274, 9548–9555 (1999).

98. Dong, A., Meyer, J. D., Brown, J. L., Manning, M. C. & Carpenter, J. F. Comparative Fourier Transform Infrared and Circular Dichroism Spectroscopic Analysis of α1-Proteinase Inhibitor and Ovalbumin in Aqueous Solution. Arch. Biochem. Biophys. 383, 148–155 (2000).

99. Mast, A. E., Enghild, J. J., Pizzo, S. V. & Salvesen, G. Analysis of the plasma elimination kinetics and conformational stabilities of native, proteinase-complexed, and reactive site cleaved serpins: comparison of alpha 1-proteinase inhibitor, alpha 1-antichymotrypsin, antithrombin III, alpha 2-antiplasmin, angiotensinogen, and ovalbumin. Biochemistry 30, 1723–1730 (1991).

100. Maas, C. & de Maat, S. Therapeutic SERPINs: Improving on Nature. Front. Cardiovasc. Med. 8, (2021).

101. Wells, M. J., Sheffield, W. P. & Blajchman, M. A. The clearance of thrombin-antithrombin and related serpin-enzyme complexes from the circulation: role of various hepatocyte receptors. Thromb. Haemost. 81, 325–337 (1999).

102. Mast, A. E., Enghild, J. J., Pizzo, S. V. & Salvesen, G. Analysis of the plasma elimination kinetics and conformational stabilities of native, proteinase-complexed and reactive site cleaved serpins: comparison of alpha-1-proteinase inhibitor, alpha-1-antichymotrypsin, antithrombin III, alpha-2-antiplasmin, angiotensinogen, and ovalbumin. Biochemistry 30, 1723–1730 (1991).

103. Hill, L. A., Bodnar, T. S., Weinberg, J. & Hammond, G. L. Corticosteroid-Binding Globulin is a biomarker of inflammation onset and severity in female rats. J. Endocrinol. 230, 215–225 (2016).

104. Mast, A. E. et al. Kinetics and physiologic relevance of the inactivation of alpha 1-proteinase inhibitor, alpha 1-antichymotrypsin, and antithrombin III by matrix metalloproteinases-1 (tissue collagenase), −2 (72-kDa gelatinase/type IV collagenase), and - 3 (stromelysin). J. Biol. Chem. 266, 15810–15816 (1991).

105. Lewis, J. G. & Elder, P. A. Corticosteroid-binding globulin reactive centre loop antibodies recognise only the intact natured protein: Elastase cleaved and uncleaved CBG may coexist in circulation. J. Steroid Biochem. Mol. Biol. 127, 289–294 (2011).

106. Grover, S. P. & Mackman, N. Anticoagulant SERPINs: Endogenous Regulators of Hemostasis and Thrombosis. Front. Cardiovasc. Med. 9, (2022).

107. Lucas, A. R. et al. Virus-derived serpin reduces immuno-coagulopathic damage in murine colitis by targeting the urokinase-type plasminogen activator receptor (uPAR) and complement. Sci. Rep. 16, 2767 (2025).

108. Rau, J. C., Beaulieu, L. M., Huntington, J. A. & Church, F. C. Serpins in thrombosis, hemostasis and fibrinolysis. J. Thromb. Haemost. 5, 102–115 (2007).

109. Park, D. J. et al. Serpin-loaded extracellular vesicles promote tissue repair in a mouse model of impaired wound healing. J. Nanobiotechnology 20, 474 (2022).

110. Simone, T. M. & Higgins, P. J. Inhibition of SERPINE1 Function Attenuates Wound Closure in Response to Tissue Injury: A Role for PAI-1 in Re-Epithelialization and Granulation Tissue Formation. J. Dev. Biol. 3, 11–24 (2015).

111. Festoff, B. W., Reddy, R. B., Vanbecelaere, M., Smirnova, I. & Chao, J. Activation of serpins and their cognate proteases in muscle after crush injury. J. Cell. Physiol. 159, 11–18 (1994).

112. Fu, Z., Thorpe, M., Akula, S., Chahal, G. & Hellman, L. T. Extended Cleavage Specificity of Human Neutrophil Elastase, Human Proteinase 3, and Their Distant Ortholog Clawed Frog PR3—Three Elastases With Similar Primary but Different Extended Specificities and Stability. Front. Immunol. 9, (2018).

113. Cooley, J., Takayama, T. K., Shapiro, S. D., Schechter, N. M. & Remold-O’Donnell, E. The Serpin MNEI Inhibits Elastase-like and Chymotrypsin-like Serine Proteases through Efficient Reactions at Two Active Sites. Biochemistry 40, 15762–15770 (2001).

114. Wang, Y., Lu, X., Wang, Z. & Zheng, W. Heat transport across multi-layered skin tissue experiencing short-pulse laser irradiation: Case of temperature-dependent thermal physical parameters. Int. J. Heat Mass Transf. 213, 124335 (2023).

115. Luchakov, Y. I. & Shabanov, P. D. Transport of heat through the skin. Rev. Clin. Pharmacol. Drug Ther. 15, 68–71 (2017).

116. Douglas, R. H. et al. Dragon fish see using chlorophyll. Nature 393, 423–424 (1998).

117. Isayama, T. et al. An accessory chromophore in red vision. Nature 443, 649–649 (2006).

118. Washington, I. et al. Chlorophyll derivatives as visual pigments for super vision in the red. Photochem. Photobiol. Sci. 6, 775–779 (2007).

119. Swenson, E. S., Price, J. G., Brazelton, T. & Krause, D. S. Limitations of green fluorescent protein as a cell lineage marker. Stem Cells 25, 2593–2600 (2007).

120. Hang, Y., Boryczka, J. & Wu, N. Visible-Light and Near-Infrared Fluorescence and Surface-Enhanced Raman Scattering Point-of-Care Sensing and Bio-imaging: A Review. Chem. Soc. Rev. 51, 329–375 (2022).

121. Lei, Z., et al. Bright, Stable, and Biocompatible Organic Fluorophores Absorbing/Emitting in the Deep Near-Infrared Spectral Region. Angew. Chem. Int. Ed. 56, 2979–2983 (2017).

122. Wegiel, B. et al. Cell Surface Biliverdin Reductase Mediates Biliverdin-induced Anti-inflammatory Effects via Phosphatidylinositol 3-Kinase and Akt*. J. Biol. Chem. 284, 21369–21378 (2009).

123. Fathi, P. et al. Biodegradable Biliverdin Nanoparticles for Efficient Photoacoustic Imaging. ACS Nano 13, 7690–7704 (2019).

124. Fu, Q., Zhu, R., Song, J., Yang, H. & Chen, X. Photoacoustic Imaging: Contrast Agents and Their Biomedical Applications. Adv. Mater. 31, e1805875 (2019).

125. Weber, J., Beard, P. C. & Bohndiek, S. E. Contrast agents for molecular photoacoustic imaging. Nat. Methods 13, 639–650 (2016).

126. Yao, J. & Wang, L. V. Recent progress in photoacoustic molecular imaging. Curr. Opin. Chem. Biol. 45, 104–112 (2018).

127. Yao, J. & Wang, L. V. Sensitivity of photoacoustic microscopy. Photoacoustics 2, 87–101 (2014).

128. Sridharan, B. & Lim, H. G. Advances in photoacoustic imaging aided by nano contrast agents: special focus on role of lymphatic system imaging for cancer theranostics. J. Nanobiotechnology 21, 437 (2023).

129. Bolger, A. M., Lohse, M. & Usadel, B. Trimmomatic: a flexible trimmer for Illumina sequence data. Bioinformatics 30, 2114–2120 (2014).

130. Haas, B. J. et al. De novo transcript sequence reconstruction from RNA-Seq: reference generation and analysis with Trinity. Nat. Protoc. 8, 10.1038/nprot.2013.084 (2013).

131. Xin, L. et al. A streamlined platform for analyzing tera-scale DDA and DIA mass spectrometry data enables highly sensitive immunopeptidomics. Nat. Commun. 13, 3108 (2022).

132. Floden, E. W. et al. PSI/TM-Coffee: a web server for fast and accurate multiple sequence alignments of regular and transmembrane proteins using homology extension on reduced databases. Nucleic Acids Res. 44, W339–343 (2016).

133. Procter, J. B. et al. Alignment of Biological Sequences with Jalview. Methods Mol. Biol. 2231, 203–224 (2021).

134. Stothard, P. The sequence manipulation suite: JavaScript programs for analyzing and formatting protein and DNA sequences. BioTechniques 28, 1102, 1104 (2000).

135. Abramson, J. et al. Accurate structure prediction of biomolecular interactions with AlphaFold 3. Nature 630, 493–500 (2024).

136. Matlashov, M. E. et al. A set of monomeric near-infrared fluorescent proteins for multicolor imaging across scales. Nat. Commun. 11, 239 (2020).

137. Pace, C. N., Vajdos, F., Fee, L., Grimsley, G. & Gray, T. How to measure and predict the molar absorption coefficient of a protein. Protein Sci. 4, 2411–2423 (1995).

138. Yu, L., Nina-Paravecino, F., Kaeli, D. R. & Fang, Q. Scalable and massively parallel Monte Carlo photon transport simulations for heterogeneous computing platforms. J. Biomed. Opt. 23, 010504 (2018).

139. Fang, Q. & Boas, D. A. Monte Carlo Simulation of Photon Migration in 3D Turbid Media Accelerated by Graphics Processing Units. Opt. Express 17, 20178–20190 (2009).

140. Prahl, S. A., Gemert, M. J. C. van & Welch, A. J. Determining the optical properties of turbid media by using the adding–doubling method. Appl. Opt. 32, 559–568 (1993).

141. Menozzi, L. et al. Three-dimensional diffractive acoustic tomography. Nat. Commun. 16, 1149 (2025).

