## Supplementary Figures for "Serpin-Driven Green Camouflage and NIR Fluorescence in Frogs"

**Supplementary Figures 1–7**

**Supplementary Video 1**

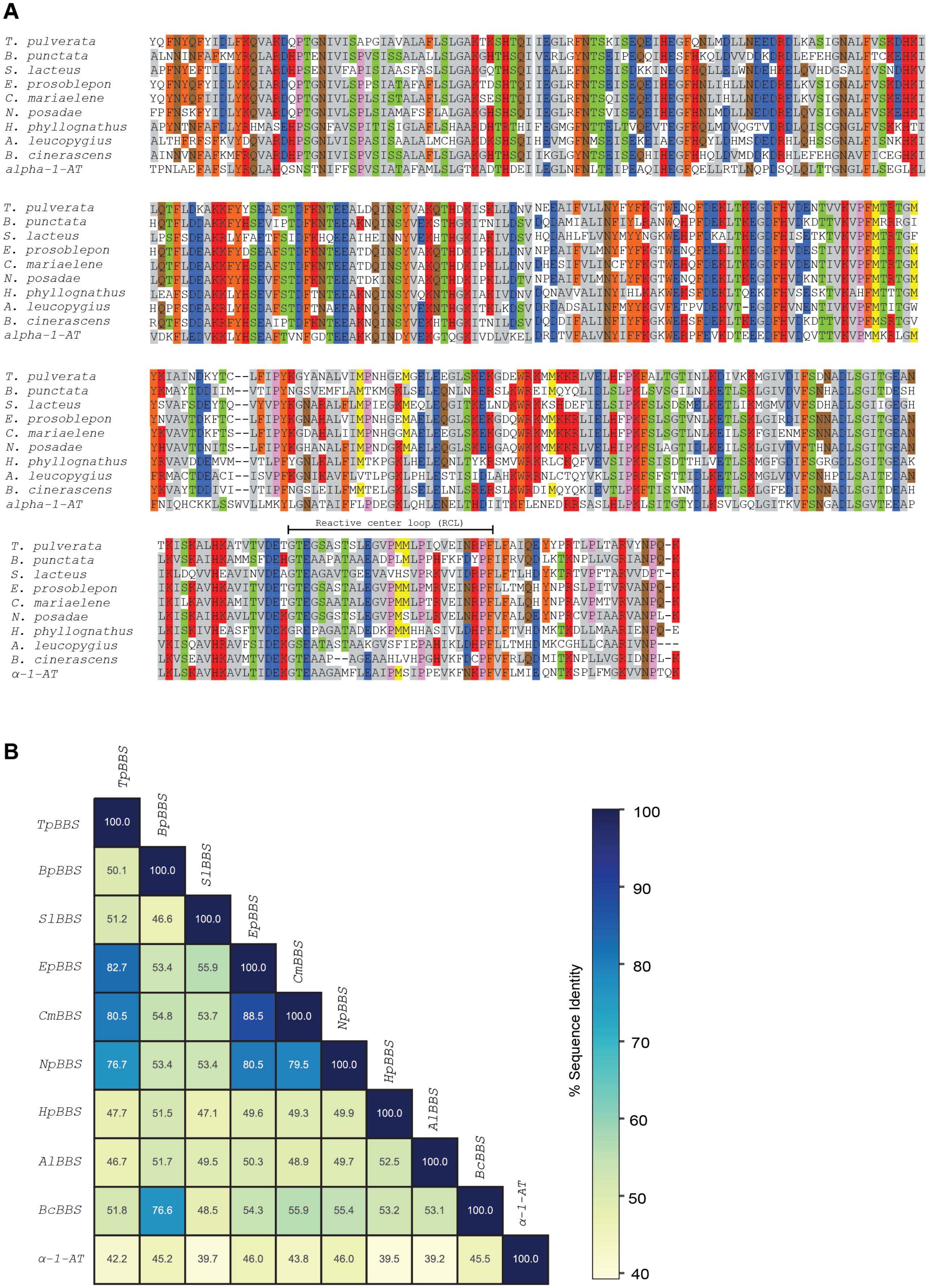
A
T. pulverata
B. punctata
S. lacteus
E. prosoblepon
C. mariaelene
N. posadae
H. phyllognathus
A. leucopygius
B. cinerascens
alpha-1-AT
Reactive center loop (RCL)
α-1-AT
B
TpBBS
BpBBS
SlBBS
EpBBS
CmBBS
NpBBS
HpBBS
AlBBS
BcBBS
α-1-AT
100.0
50.1 100.0
51.2 46.6 100.0
82.7 53.4 55.9 100.0
80.5 54.8 53.7 88.5 100.0
76.7 53.4 53.4 80.5 79.5 100.0
47.7 51.5 47.1 49.6 49.3 49.9 100.0
46.7 51.7 49.5 50.3 48.9 49.7 52.5 100.0
51.8 76.6 48.5 54.3 55.9 55.4 53.2 53.1 100.0
42.2 45.2 39.7 46.0 43.8 46.0 39.5 39.2 45.5 100.0
% Sequence Identity
100
90
80
70
60
50
40

**Supp. Fig. 1. (A)** Multiple sequence alignment of various BBSs from different glassfrogs and treefrogs along with human alpha-1-antitrypsin. N-terminus region of the protein was not included in the alignment. **(B)** Sequence identity matrix generated from the multiple sequence alignment.

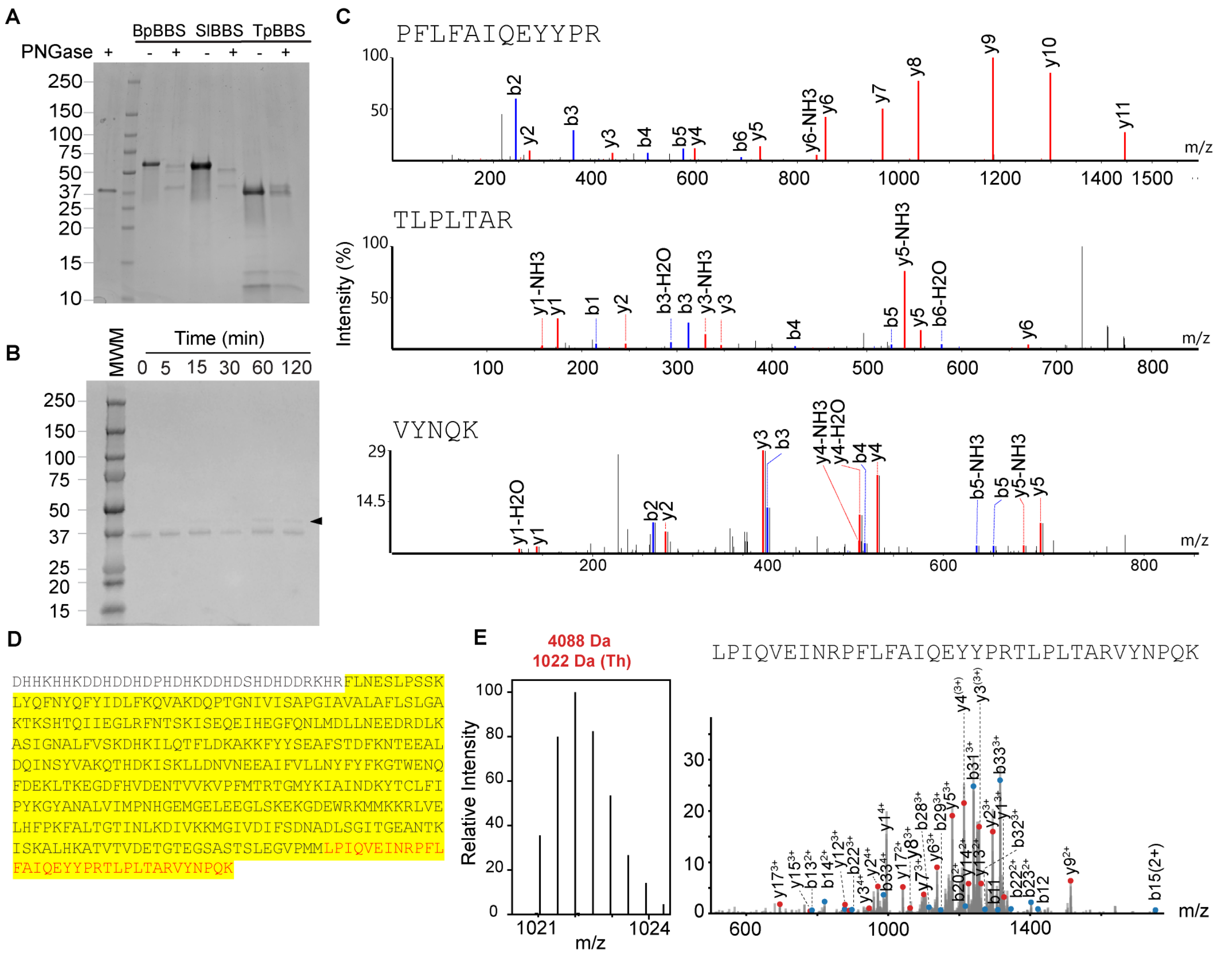


**Supp. Fig. 2. (A)** Glycosylation in BBS. SDS PAGE gel showing mobility differences between PNGase F-treated and untreated BBS. Treatment with PNGaseF revealed two bands for BpBBS and one for SlBBS, consistent with the number of glycosylation sites inferred from their sequences. TpBBS did not exhibit any changes between the treated and untreated conditions. **(B)** BS3 *d0-d4* crosslinking of endogenous TpBBS from 0-120 minutes incubation showed a band with a larger molecular weight (black arrow) –compatible with the addition of the C terminus– appearing after 15 minutes. **(C)** Bottom-up proteomics (in-gel tryptic digestion) of the crosslinked TpBBS from the gel in (B). Analyses of LC-MS/MS revealed multiple fragments within the C-terminus of TpBBS. **(D)** The coverage of the bottom-up mass spectrometry analysis of TpBBS –highlighted in yellow–increases from 80% to 91% when the protein is crosslinked. C-terminus of TpBBS after the scissile bond is shown in red. **(E)** LC-MS/MS of endogenous TpBBS revealed a fragment of

4,088 Da aligning which aligns with the predicted molecular weight of the C-terminus of the protein. MS (left) and MS/MS of 1022 m/z (4+) peak (right).

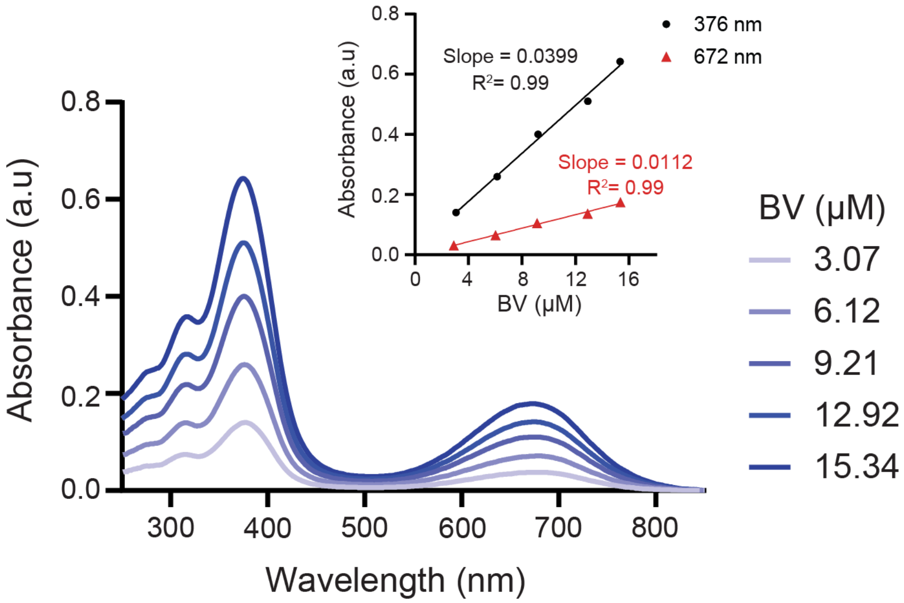


**Supp. Fig. 3. Molar extinction coefficient of biliverdin in aqueous solution (1X PBS).** Absorption spectrum of various concentrations of biliverdin in 1X PBS. *Inset*, Absorbance at 376 and 672 nm were plotted against the increasing concentrations of biliverdin. The slopes (molar absorption coefficients) were determined from the linear regression of the plotted values as $\varepsilon_{376\ nm} = 39{,}900\ M^{-1}cm^{-1}$ and $\varepsilon_{672\ nm}=11{,}200\ M^{-1}cm^{-1}$.

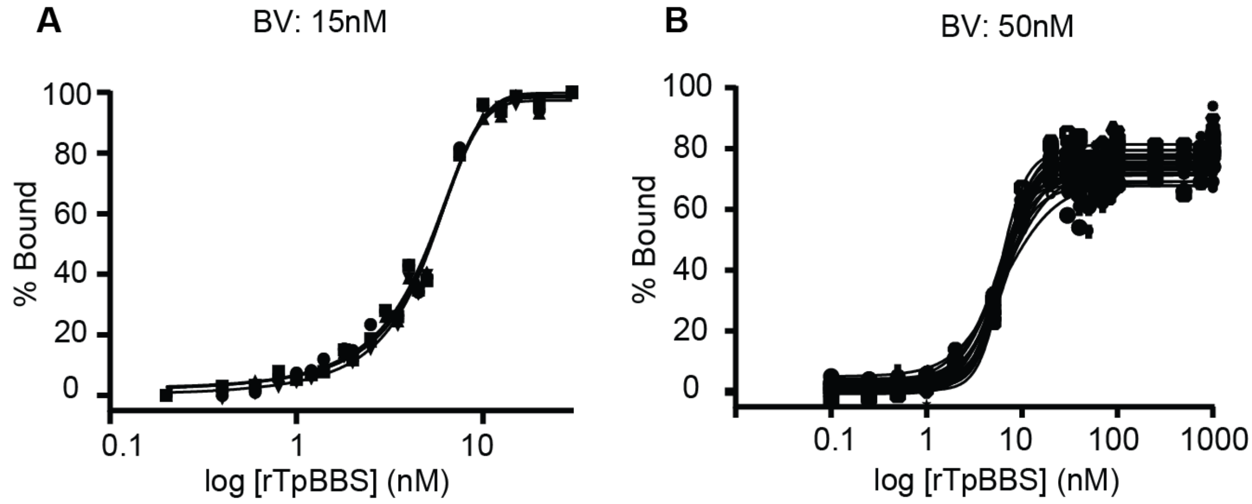


**Supp. Fig. 4. Binding regime of rTpBBS.** The fluorescence of biliverdin (BV) bound to rTpBBS remained stable over time, indicating that binding equilibrium was reached. This was observed in (A) with 15 nM BV over 0–4 h and (B) with 50 nM BV over 0–24 h. The average apparent $K_d$ values calculated from these curves were 5 nM (A) and 6.2 nM (B). Further lowering the BV concentration reduced fluorescence below the detection limit (data not shown).

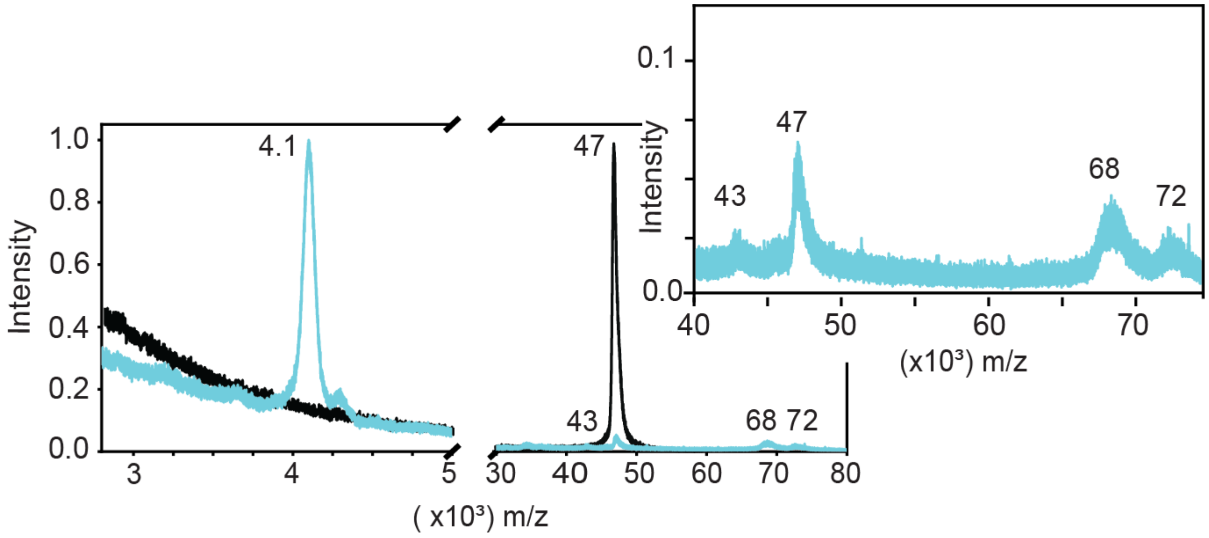


**Supp. Fig. 5. MALDI-TOF of rTpBBS in relaxed- (cyan) and stressed- (black) conformation.** The appearance of the ~4.1 kDa signal indicates the cleavage of rTpBBS by human neutrophil elastase, a fragment not observed in untreated rTpBBS. The 43 kDa and 47 kDa peaks correspond to the cleaved (relaxed) serpin fold and native (stressed) rTpBBS, respectively. The 68 kDa and 72 kDa peaks correspond to elastase complexed with the cleaved rTpBBS and native rTpBBS, respectively. The molecular weight of neutrophil elastase is ~26 kDa.

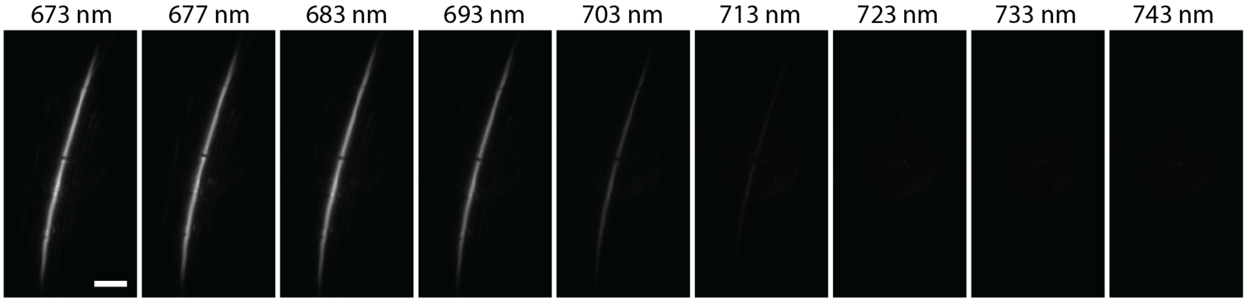


**Supp. Fig. 6. PA images of BBS-filled tubes for spectral determination.** TpBBS is filled into a silicone tube with an inner diameter of 0.31 mm and an outer diameter of 0.64 mm. Then, PA images are acquired at incremental wavelengths from 673 to 743 nm. The averaged PA amplitude of the tube at each wavelength is used for the determination of the PA spectra in **Fig. 6**. Scale bar: 5 mm.

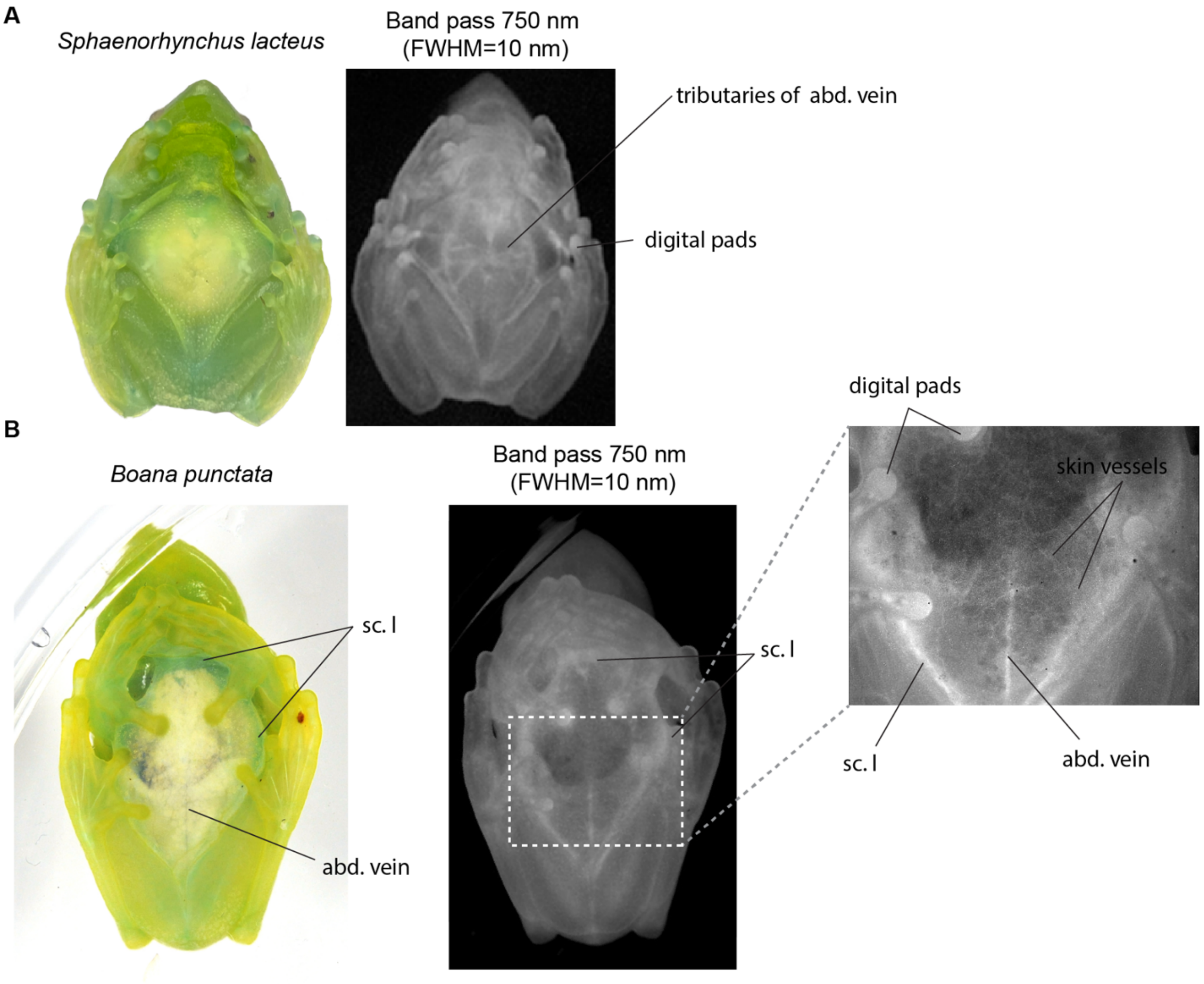


**Supp. Fig. 7. BBS-mediated NIR fluorescence in *Sphaenorhynchus lacteus* and *Boana punctata***. The brightfield images were shown alongside the fluorescent images to visualize the internal tissues. *Inset,* enlarged portion of the ventral side of *B. punctata* highlights the anatomical structures**.** *ab. v.*= abdominal vein; *sc. l.* = subcutaneous lymph

**Supp. Video 1**

Ventral view of a glassfrog (*T. pulverata*) under white light illumination (left) and under 660 nm light to excite BBS fluorescence emission (750 nm bandpass filter, FWHM = 10 nm). Frog NIR fluorescence is shown in grayscale (center) or as orange pseudocolor (right). BBS is distributed in the vasculature, lymph, bones, and muscles.
